# Characterization of systemic genomic instability in budding yeast

**DOI:** 10.1101/2020.05.25.115535

**Authors:** Nadia M. V. Sampaio, V. P. Ajith, Ruth A. Watson, Lydia R. Heasley, Parijat Chakraborty, Aline Rodrigues-Prause, Ewa P. Malc, Piotr A. Mieczkowski, Koodali T. Nishant, Juan Lucas Argueso

**Affiliations:** Department of Environmental and Radiological Health Sciences, Colorado State University, Fort Collins-CO 80523, USA; Cell and Molecular Biology Graduate Program, Colorado State University, Fort Collins-CO 80523, USA; School of Biology, Indian Institute of Science Education and Research, Thiruvananthapuram, Trivandrum 695016, India; Department of Genetics, University of North Carolina, Chapel Hill-NC 27599, USA

## Abstract

Conventional models of genome evolution are centered around the principle that mutations form independently of each other and build up slowly over time. We characterized the occurrence of bursts of genome-wide loss-of-heterozygosity (LOH) in *Saccharomyces cerevisiae*, providing support for an additional non-independent and faster mode of mutation accumulation. We initially characterized a yeast clone isolated for carrying an LOH event at a specific chromosome site, and surprisingly, found that it also carried multiple unselected rearrangements elsewhere in its genome. Whole genome analysis of over 100 additional clones selected for carrying primary LOH tracts revealed that they too contained unselected structural alterations more often than control clones obtained without any selection. We also measured the rates of coincident LOH at two different chromosomes and found that double LOH formed at rates 14-150 fold higher than expected if the two underlying single LOH events occurred independently of each other. These results were consistent across different strain backgrounds, and in mutants incapable of entering meiosis. Our results indicate that a subset of mitotic cells within a population can experience discrete episodes of systemic genomic instability, when the entire genome becomes vulnerable and multiple chromosomal alterations can form over a narrow time window. They are reminiscent of early reports from the classic yeast genetics literature, as well as recent studies in humans, both in the cancer and genomic disorder contexts. The experimental model we describe provides a system to further dissect the fundamental biological processes responsible for punctuated bursts of structural genomic variation.

**SIGNIFICANCE STATEMENT:** Mutations are generally thought to accumulate independently and gradually over many generations. Here, we combined complementary experimental approaches in budding yeast to track the appearance of chromosomal changes resulting in loss-of-heterozygosity (LOH). In contrast to the prevailing model, our results provide evidence for the existence of a path for non-independent accumulation of multiple chromosomal alteration events over few generations. These results are analogous to recent reports of bursts of genomic instability in human cells. The experimental model we describe provides a system to further dissect the fundamental biological processes underlying such punctuated bursts of mutation accumulation.

## INTRODUCTION

Alterations in the DNA sequence and structure of chromosomes are driving forces in evolution and disease. Thus, understanding the mechanisms and dynamics through which mutations form and accumulate in genomes has important implications for wide-ranging biological processes, such as microbial adaptation to environmental changes, as well as cancer onset and progression. The conventional paradigm for genome evolution postulates that new mutations occur randomly and independently of each other, and accumulate gradually over time as organisms develop and species diverge. While this primary path is well established and robustly supported, evidence also exists to suggest a substantial contribution from additional processes that lead to rapid, punctuated mutation accumulation, particularly in the cancer context (1, 2).

Recent studies of intratumoral genomic heterogeneity support the occurrence of transient bursts of structural chromosomal instability leading to the formation of multiple copy number alterations (CNAs) over short time periods (3–6). A similar pattern of rapid CNA accumulation followed by stable clonal propagation has also been reported in the human germline and during early embryonic development leading to genomic disorders (7). Interestingly, these findings are broadly analogous to accounts of frequent detection of coincident mitotic recombination and other rare types of chromosomal alterations recorded in the classic yeast genetics literature (8–13), and also described in contemporary studies (14–17).

We leverage the budding yeast *Saccharomyces cerevisiae* to track the emergence of coincident chromosomal changes using high resolution genomic approaches. We showed that yeast clones selected for carrying specific and relatively rare loss-of-heterozygosity (LOH) events often contained additional unselected alterations in other chromosomes. We quantified the magnitude of this non-independent accumulation of structural variation using fluctuation assays, and characterized it qualitatively through genome-wide mapping of recombination tracts. Our results suggested that, in addition to the established independent and gradual mutation accumulation path, wild type yeast cells can also experience transient recombinogenic states when instability affects the entire genome systemically, leading to a punctuated pattern of LOH and CNA accrual. The experimental approach described here provides a tractable model system to further dissect the fundamental mechanisms underlying systemic genomic instability (SGI) processes, including those that may play a role in human disease development.

## RESULTS

### Abundant unselected genomic alterations in a spontaneous yeast isolate

Our group recently reported a case study of mitotic recombination leading to a readily discernable colony morphology alteration in the *S. cerevisiae* diploid strain JAY270 (18). We showed that this strain is heterozygous for a frameshift mutation in *ACE2* (*ACE2/ace2-A7*), a transcriptional regulator of genes involved in septum destruction following cytokinesis (19). Clonally derived cells carrying an LOH tract spanning the region of the right arm of chromosome XII containing the *ACE2* locus (Chr12, homozygosity for the maternal [M] haplotype; *ace2-A7/ace2-A7*) acquired a mother-daughter cell separation defect that manifested macroscopically as a distinctive rough colony morphology phenotype in agar media (Fig. 1B inset). The presence of heterozygous single nucleotide polymorphisms (HetSNPs) associated with restriction enzyme recognition sites allowed us to characterize the LOH tracts spanning the *ACE2* locus in five independent rough clones. In all cases, these tracts were consistent with allelic interhomolog mitotic recombination leading to segmental copy-neutral LOH (20). Unexpectedly, in one of the rough isolates (JAY664), our analysis revealed the presence of an additional unselected LOH tract on the left arm of Chr12 leading to homozygosity for the paternal haplotype (Chr12P). Based on the measured rates of LOH in JAY270, we estimated that these two independent tracts should have coincided in the same clone at a very low rate, in the ∼10^-10^ to 10^-11^/cell/division range.

**Figure 1.**
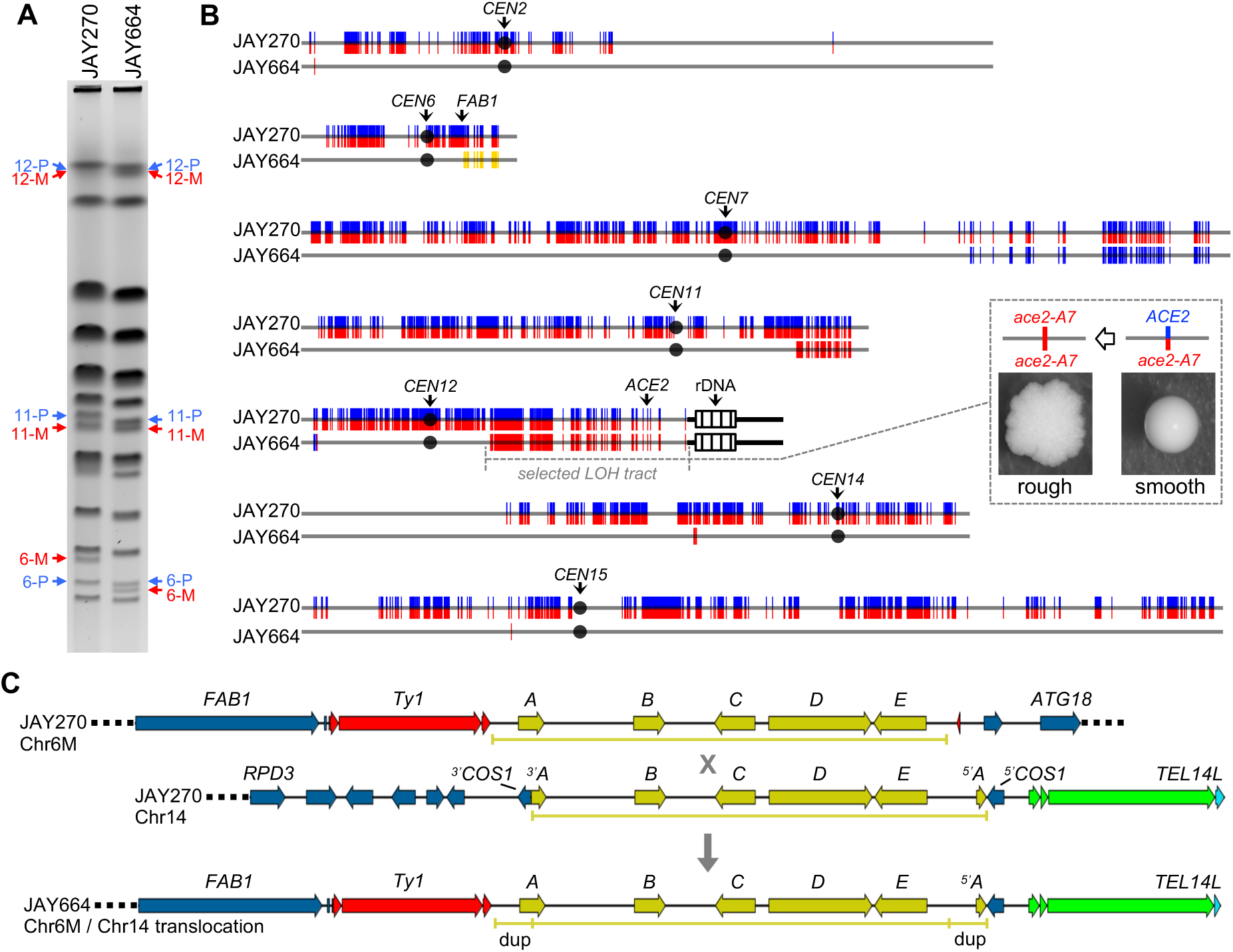
Genomic analysis of the JAY664 rough colony isolate. **A** PFGE of JAY270 and JAY664. Arrows indicate the bands for Chr6, Chr11 and Chr12, with the two homologs identified as paternal (P, blue) or maternal (M, red) according to the phased HetSNP haplotypes described previously (18). **B** Distribution of HetSNPs in JAY270 and tracts of LOH and CNA in JAY664. JAY270 HetSNPs are represented as double colored vertical lines (P/M; blue/red). In the JAY664 maps below, markers that remained heterozygous were omitted to emphasize visualization of tracts of copy-neutral LOH (double red [M/M] and double blue [P/P]), or CNA deletion on Chr6 (double yellow). The LOH tract on the right arm of Chr12 spanning the *ACE2* locus was selected based on the rough colony morphology shown in the inset. The LOH tracts on the left arm of Chr12 and on each of the other chromosomes were unselected. Chromosome numbers are to the left of each map, black circles represent their respective centromeres, and the positions of pertinent loci are indicated. Striped boxes on the right arm of Chr12 represent ∼1.5 Mb of ribosomal DNA repeats (rDNA); regions distal to the rDNA are homozygous in JAY270 so are not represented. Chromosome plots were generated to scale in Python 2.7 using the matplotlib package and a custom script. For size reference, Chr6 is 270 kb. **C** shows the detailed DNA structures determined by Nanopore single molecule long read WGS present in JAY270 Chr6M, JAY270 Chr14, and at the junction of the JAY664 Chr6M/Chr14 NAHR-mediated non-reciprocal translocation. The sequences in yellow correspond to the *Zb* circle insertions, with the five ORFs being labeled *A* through *E* following the abbreviations described previously (21). The X represents the deduced NAHR event and the downward arrow points to the recombination outcome detected in JAY664. Ty1 and delta LTR retrotransposon sequences are shown in red, and telomeric sequences (X and Y’ elements, and the telomeric short repeats) are shown in green. *dup* indicates the short segment of the *Zb* circle at the Chr6-M / Chr14 junction that was duplicated by the NAHR event.

Sparse cases of mitotic recombination events coinciding at two marked loci have been previously reported in yeast species (8, 10–16). We considered that a similar process may have been at play in the formation of JAY664 isolate, in which case other parts of its genome might also have been affected. Thus, we carried out a full characterization of structural genomic variation in this clone, initially using pulse-field gel electrophoresis (PFGE), followed by two independent and complementary whole genome sequencing (WGS) platforms. PFGE detected the presence of at least two unselected gross rearrangements in chromosomes other than Chr12 (Chr11P and Chr6M; Fig. 1A). Next, analysis of Illumina short read WGS data revealed that the JAY664 genome contained a remarkable total of seven unselected structural changes, in addition to the selected LOH tract spanning the *ACE2* locus (Fig. 1B, Table S3, and File S1). These included the terminal copy-neutral LOH tract (P/P) on the left arm of Chr12 identified previously (18), and two terminal LOH tracts associated with the rearrangements detected by PFGE. Chr11 contained a terminal copy-neutral LOH tract (M/M) on the right arm, and Chr6M had a terminal CNA deletion tract on the right arm distal to *FAB1* (P/-; only P HetSNPs detected at ∼50% relative read depth coverage). Additional LOH tracts that did not result in altered copy number or chromosome length were identified on Chr7 (terminal), Chr2, Chr14 and Chr15 (interstitial).

We also generated Oxford Nanopore single molecule long WGS reads from JAY270 and JAY664 to characterize additional structural features near the tract endpoints detected by short read WGS, and thus infer the respective causal recombination mechanisms. We identified sets of long Nanopore reads spanning the entirety of each of the three JAY664 interstitial LOH tracts (>41.4 kb for the Chr2 LOH tract, and ∼10 kb for the Chr14 and Chr15 tracts) and aligned them to determine the phasing configuration of the flanking HetSNP markers (Fig. S1A). This analysis validated all homozygous base calls made at these positions using short read WGS data and showed that there was no alteration of the JAY270 parental phasing of the HetSNPs flanking the LOH regions. Therefore, all three unselected interstitial LOH tracts detected in JAY664 were formed through an interhomolog gene conversion (GC) mechanism without associated reciprocal crossovers (20).

Next we focused on the junction associated with the Chr6M deletion by identifying long reads (>40 kb) that included *FAB1* (Fig. 1C). In the JAY270 parent strain we identified two discrete groups of read structures corresponding to the Chr6P and Chr6M homologs with their respective phased HetSNPs. This region in JAY270 Chr6P was similar in structure to the S288c reference (data not shown), but JAY270 Chr6M contained an insertion that included a full-length Ty1 element and a ∼17 kb DNA segment absent in S288c (Fig. S1B). This segment corresponded to a well-characterized set of five ORFs acquired by *S. cerevisiae* through horizontal gene transfer of an extrachromosomal circular DNA intermediate postulated to have originated in *Zygosacchromyces bailii* (*Zb* circle) (21, 22). A similar analysis of Nanopore data derived from JAY664 revealed that Chr6M reads included on their distal side ∼9 kb of sequences matching the left end of S288c Chr14, from a position in the middle of the *COS1* gene to the telomeric repeats (*TEL14L*). This prompted us to retrieve long reads from JAY270 and JAY664 corresponding to the Chr14 left arm region spanning from *RPD3* to *TEL14L*. This region, which is homozygous in JAY270, contained another insertion of the *Zb* circle within *COS1* (Fig. S1C), thus offering a substantial region of sequence homology to Chr6M. Notably, these two *Zb* circle insertions had different arrangement of the five ORFs, consistent with the circle linearizing prior to integration between ORFs *A* and *E* in Chr6M, and within ORF *A* in Chr14. This difference was reflected in the structure of the JAY664 Chr6-M/Chr14 translocation junction, which included a ∼1.5 kb flanking duplication spanning the promoter and 5’ region of ORF *A*. This indicated that the non-allelic homologous recombination (NAHR; (20)) event responsible for the formation of the translocation occurred somewhere in the 15.5 kb region between the 3’ end of ORF *A* and the promoter of ORF *E*.

### Characterization of coincident recombination in JAY270

The remarkable number of unselected rearrangements detected in the JAY664 genome made it unlikely that they had all accumulated independently and over time. A simpler explanation would be that instead, this highly altered karyotype formed rapidly during a discrete burst of structural variation in one or a few ancestor cells in the JAY664 lineage. If such an underlying process, which here we refer to as systemic genomic instability (SGI), does indeed exist and is active in *S. cerevisiae*, then selecting for clones carrying a primary LOH tract should enrich for the presence of unselected LOH tracts or other forms of structural variation elsewhere in the genome. In the specific case of JAY270, rough colony derivatives selected for carrying an LOH tract on Chr12 should carry more unselected tracts than comparable control smooth clones isolated without any selection.

To test this prediction, we obtained 28 independent unselected smooth control clones, and screened additional cultures until 23 independent rough morphology clones were found (28 total rough clones including the five isolated previously (18)). The controls were isolated by picking random smooth colonies plated after growth for five consecutive transfer cycles in liquid culture (∼57 cell generations; Methods). The experimental clones were derived from the same liquid growth regimen, but instead plating ∼1,000 cells after every passage cycle to screen for the presence of rough colonies until one clone per culture was isolated. Thus, the rough clones were isolated after a variable number of cell generations between cultures, but typically lower (median ∼50; Table S3) than the number used to isolate the controls. Initial PCR and PFGE analyses of these isolates showed that all smooth clones remained heterozygous at the *ACE2* locus and had normal karyotypes, whereas the rough clones were all homozygous for *ace2*-*A7* and seven of them contained discernible size alterations of chromosomes other than Chr12 (data not shown).

We next we carried out Illumina short read WGS analysis of these clones to characterize any differences in frequency or qualitative spectra of unselected structural variation between clone sets (Fig. 2 and Table S3). A total of 12 unselected LOH tracts were identified in 9 of the 28 smooth control clones, most of which had only one tract per clone. The rough clones collectively carried a larger and more diverse pool of unselected genomic changes. Including the JAY664 tracts described above, a total of 26 unselected structural variation tracts (24 copy-neutral LOH and 2 CNA deletions) were found in 12 of the 28 rough clones, most of which had two or more tracts per clone. The overall proportions of unselected tracts between the clone sets were statistically different (p<0.001). Further, the incidence of unselected tracts per clone among the controls followed a Poison random distribution (p=0.585), whereas among the rough clones it did not (p<0.001). These results were consistent with enhanced and non-independent accumulation of unselected LOH and CNA in clones selected for carrying a primary recombination tract on Chr12. This was particularly notable since most rough clones were isolated after a lower number of cell generations than the smooth controls, and, there was no significant correlation (p=0.305; linear regression R^2^=0.323) between the number of generations and the number of unselected tracts detected per rough clone.

**Figure 2.**
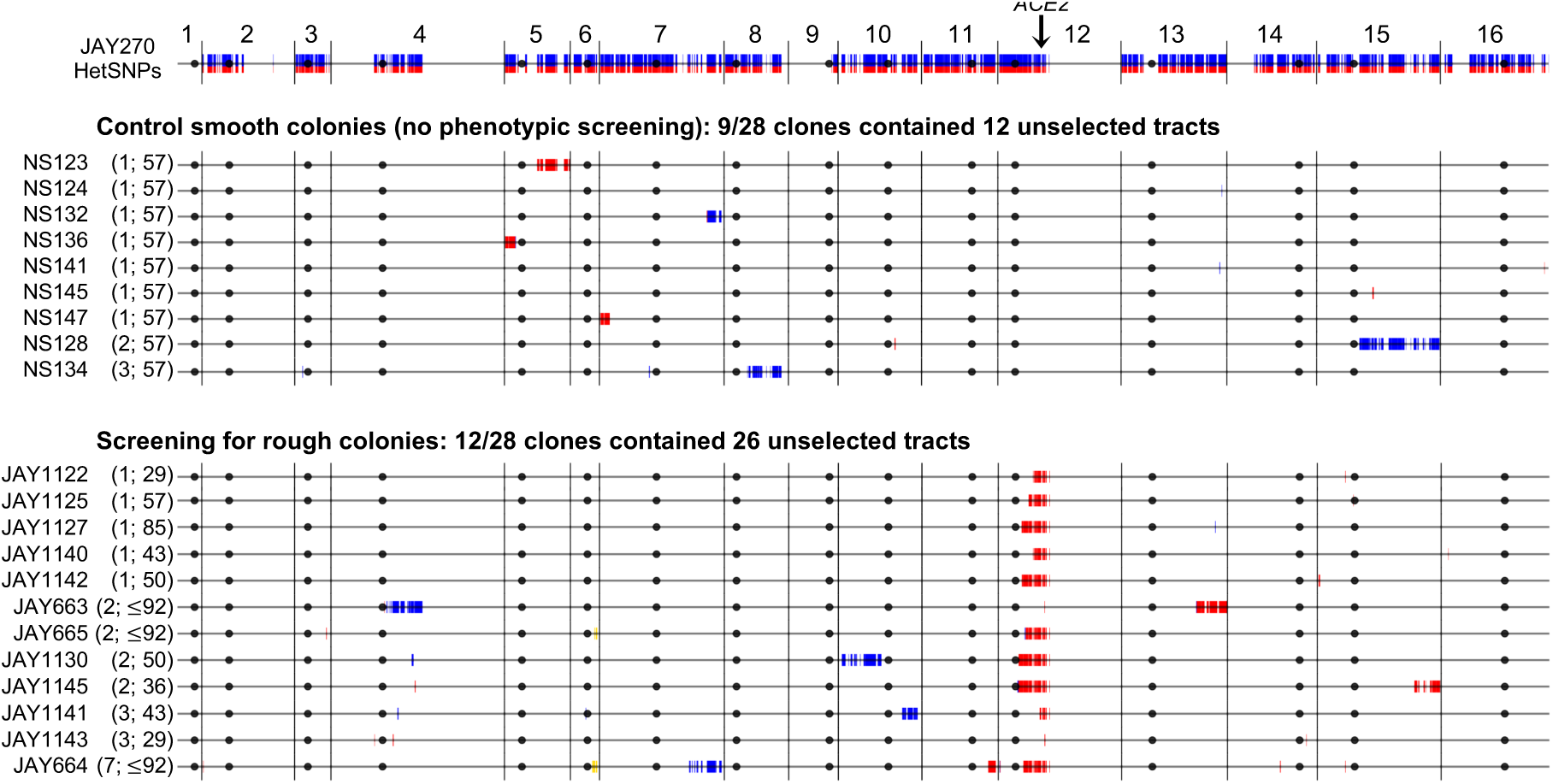
Genome-wide maps of unselected LOH and CNA tracts in smooth and rough colony isolates derived from JAY270. The top horizontal line is the linear end-to-end depiction of the 16 *S. cerevisiae* chromosomes in the JAY270 strain, with HetSNPs represented as described in Fig.1B. Chromosome numbers are indicated above, black circles represent their respective centromeres, and the position of the *ACE2* locus on Chr12 is shown. The tract maps below are grouped for control smooth and rough colony clones. Each horizontal line corresponds to the genomes of clones that displayed at least one unselected LOH or CNA tract. HetSNPs that were homozygous P/P (blue) or M/M (red) are shown, while markers that remained heterozygous in those clones were omitted. Two CNA deletions are shown in yellow (Chr6 [P/-] in JAY664 and Chr6 [M/-] JAY665). The numbers of unselected tracts and the number of generations used to isolate each clone are shown between parentheses (*i.e.*, (# unselected tracts; # generations)). As expected from selection for the rough colony morphology, all rough clones were homozygous for the maternal *ace2-A7* allele (red; close-up view in Fig S2). Full LOH calls with coordinates are available in File S1. Plots were generated to scale as in Fig. 1B. For size reference, Chr1 is 230 kb.

Interestingly, while the frequency and distribution of unselected LOH tracts were different, the qualitative spectra were similar (Table S3). The two clone sets had roughly 1:1 ratios of interstitial to terminal unselected LOH tracts, and the median tract sizes were also comparable (5.0 and 5.3 kb for interstitial, and 254 and 232 kb for terminal, in smooth and rough clones respectively). The WGS analysis also provided a detailed picture of the qualitative spectrum of selected Chr12 LOH in the rough clones (Fig. S2) that corroborated and expanded on our earlier report (18).

### Quantitative analyses of coincident LOH

In the second phase of this study, we set out to validate and more broadly characterize the SGI phenomenon by pursuing two additional orthogonal approaches. These experiments were carried out in diverse strain backgrounds, using fewer cell generations, and selecting for LOH at other regions of the *S. cerevisiae* genome.

We conducted a set of experiments to determine whether the degree of coincidence between LOH events at different genomic regions could be quantified. To do that we generated diploid strains hemizygous for the counter selectable CORE2 cassette (*KlURA3-ScURA3-KanMX4*) inserted on Chr13 or Chr4, and heterozygous *CAN1/can1Δ* on Chr5 (Fig. 3A-B). We then used these strains to measure the rates of LOH at these individual loci by selection for resistance to 5-FOA or canavanine, respectively, and also to measure the rate of coincident double LOH through selection for clones in plates containing both 5-FOA and canavanine. We reasoned that if the occurrence of LOH at a CORE2 insertion locus were completely independent from the occurrence of LOH at the *CAN1* locus, then the rate of coincident LOH at both loci should be equal to the multiplicative product of the two single rates. The rates of single LOH at each locus were measured to be in the 10^-5^ to 10^-4^ /cell/division range (Lea and Coulson method of the median (L&C); (23)), thus the rates of double LOH should be in the 10^-10^ to 10^-9^ /cell/division range. Instead, the double LOH rates that we measured experimentally were 14-50 fold higher than the expected rates calculated based on independence. This was initially observed in two different diploid strain backgrounds, JAY270 and CG379 (an S288c-related strain), for double LOH events at the two pairs of genomic regions assayed (Fig. 3A-D, Table S4).

**Figure 3.**
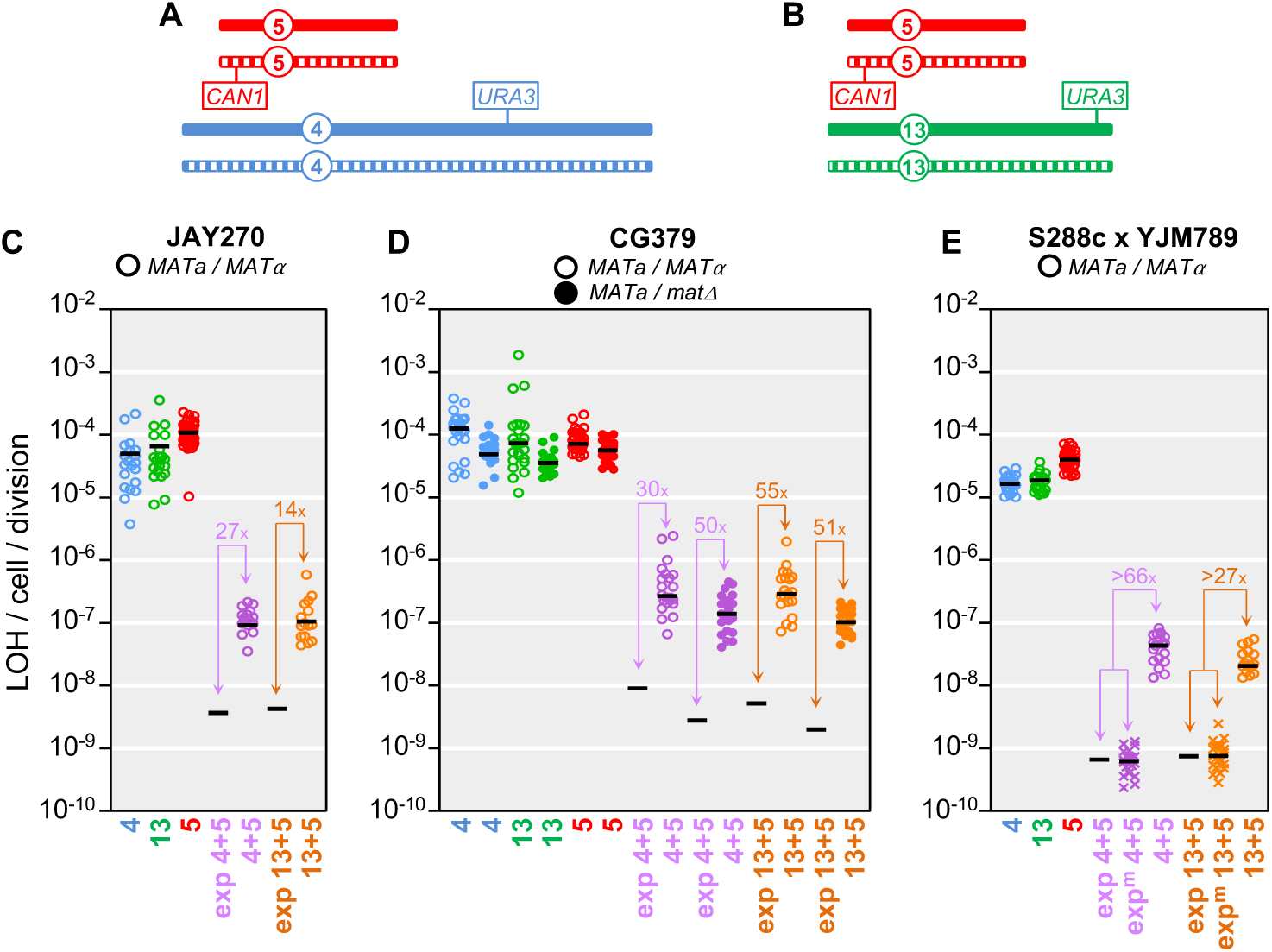
Quantitative analyses of single and double LOH rates. **A** and **B** show schematic representations of the positions of hemizygous counter-selectable markers in the diploid yeast strains used in the LOH assays. The pairs of homologous chromosomes are shown as solid or striped for Chr5 (red), Chr4 (blue), Chr13 (green), with the overall sizes, and positions of markers and centromeres drawn to approximate scale. **A** shows the genomic configuration of markers in strains used in the Chr4 plus Chr5 double LOH assays, and **B** shows the configuration used for the Chr13 plus Chr5 assays. Specifically for the hybrid strain background in **E**, the solid chromosome corresponds to the S288c homolog and the striped homolog corresponds to the YJM789 homolog. The *CAN1* locus was present at its native position on Chr5, and the CORE2 cassettes were inserted near *SSF2* in Chr4 and *ADH6* in Chr13. **C** shows the mutation rate data for the JAY270 strain background calculated using the Lea & Coulson method of the median. Circles indicate the experimentally determined rates in individual cultures, color-coded according to single chromosomes as in **A** and **B**, and double chromosomes using purple for Chr4 plus Chr5 or orange for Chr13 plus Chr5. Black horizontal bars indicate the median rate values for each culture set. The same numerical data are shown in Table S4, including 95% confidence intervals (not plotted in **C**-**E**). “exp” in the x-axis designates the expected double LOH rates based on the multiplicative product of the pertinent median single LOH rate pairs. Only the black bar is shown in these “exp” cases. The excess fold ratio of experimentally observed over calculated expected double LOH rates are shown in all pairwise comparisons. **D** shows the LOH rate data in the CG379 strain background similarly to **C**, with the exception that open circles correspond to *MATa/MATα*, while solid circles correspond to data collected in the isogenic *MATa/matΔ* strain. **E** shows the LOH rate data in the S288c x YJM789 hybrid strain background similarly to **C**, with addition of an alternative method for the calculation of expected double LOH rates (exp^m^). For this strain background, aliquots from each individual culture were plated in YPD, 5-FOA, Can, and 5-FOA plus Can (Fig. 4A), producing culture-matched single and double LOH rate results. The “X” symbols in the exp^m^ datasets correspond to the expected matched double LOH rates for each culture obtained by multiplying the pertinent two single LOH rates within that same culture. The black bar in those cases corresponds to the median value of matched expected double LOH rates. The conventional (exp) and matched expected (exp^m^) double LOH rate estimates were very close. The most conservative value for the excess ratio of observed over exp or observed over exp^m^ is shown in the plot.

We also considered the possibility that the SGI mechanism causing higher than expected rates of double LOH could result from rare initiation of meiotic recombination in a few cells in the vegetatively-grown population followed by return-to-growth (RTG) (24). Unlike the sporadic nature of mitotic recombination, meiotic recombination is initiated as a well-coordinated systemic and genome-wide process (25). To distinguish between mitotic and meiotic origins, we repeated the double LOH measurements in CG379 diploids deleted for the *MATα* locus. These *MATa/matΔ* diploids behaved essentially as *MATa* haploids. They mated efficiently to a *MATα* haploid tester strain and completely lost the ability to sporulate (data not shown). Since *MATa/matΔ* diploids are unable to activate the meiotic program, they are also unable to initiate meiotic recombination. The rate of single LOH in the *MATa/matΔ* diploids was slightly lower than the rate in the isogenic CG379 *MATa/MATα* diploids, consistently with earlier studies (26). The observed rate of double LOH in the *MATa/matΔ* strains (Fig. 3D) was also slightly lower than in *MATa/MATα*, but more importantly, it remained >51 fold higher than expected if the single LOH events were initiated independently of each other. This result indicated that the excess double LOH detected in these assays was not caused by cryptic meiotic initiation or RTG within vegetative cultures.

In order to expand our analysis to encompass the full spectrum of coincident LOH and CNA, we carried out additional experiments in a third, hybrid diploid strain background resulting from a cross between diverged haploid parents (27, 28). Two experimental strains (Fig. 3A-B) were obtained by crossing S288c *MAT*a *can1Δ* carrying a CORE2 cassette integrated on Chr4 or Chr13, to a *MATα* derivative of the clinical isolate YJM789 (*CAN1 ura3*). The resulting hybrid diploids contained ∼50,000 HetSNP markers evenly distributed across the genome that were later used to map structural variation tracts with higher resolution and sensitivity than in JAY270.

We used these new strains to quantify single and double LOH rates, initially using the same general approach described above for the JAY270 and CG379 strain backgrounds. One distinction, however, was that for these experiments we plated aliquots from each of the cultures onto permissive and all three selective media (Fig. 4A), and therefore obtained culture-matched measurements of the two single as well as the double LOH rates for each culture. This design supported two different ways of calculating the expected rate of independent double LOH: the conventional way (exp) by simply multiplying the two median single LOH rates; and an alternative method (exp^m^) by multiplying the two matched single LOH rates within each culture, and then taking the median expected double LOH rate for all cultures of that strain. These two methods produced very similar expected double LOH rate estimates (Fig. 3E), and again, were substantially lower (27-66 fold) than the experimentally measured double LOH rates.

**Figure 4.**
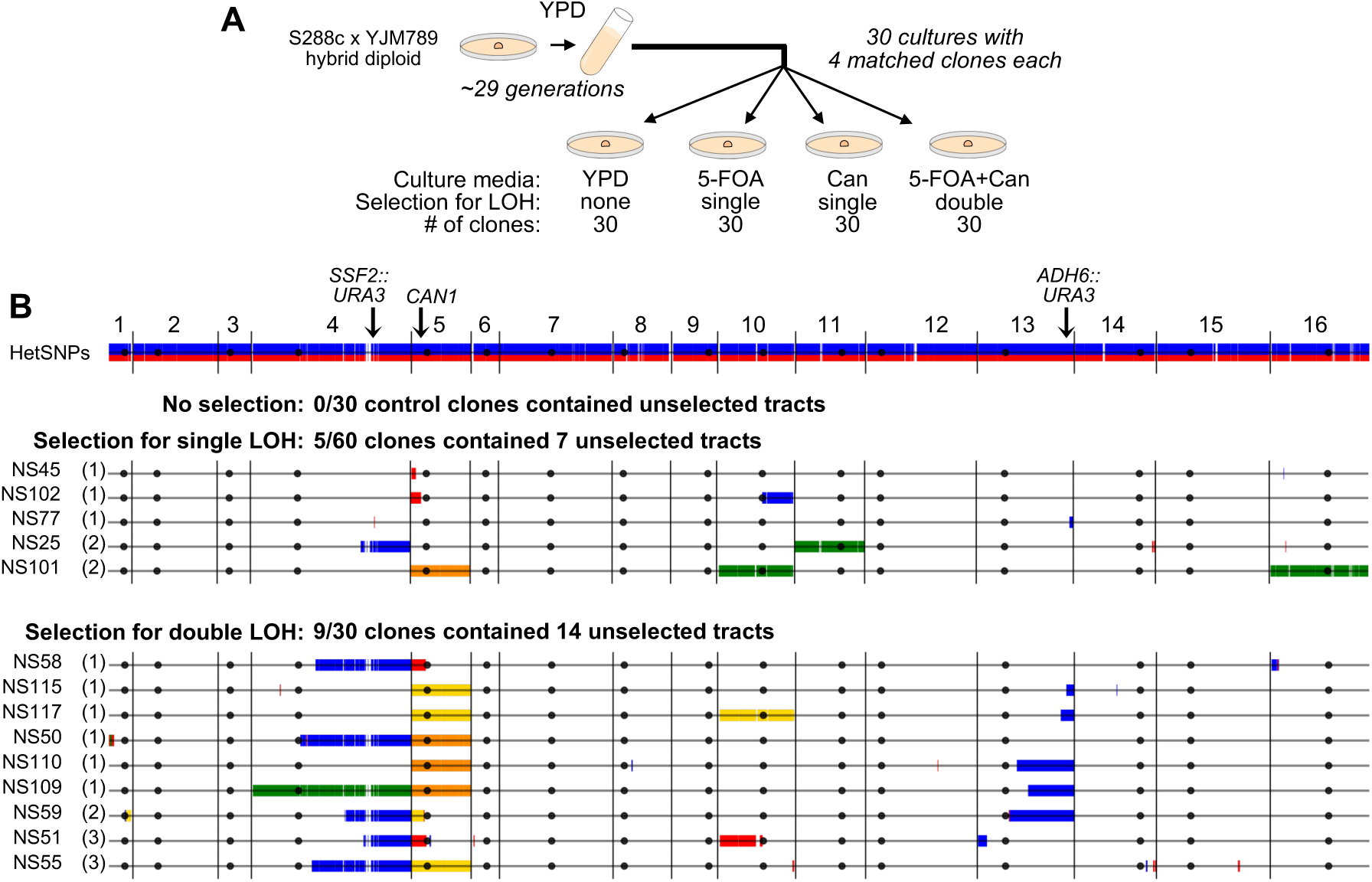
Genome-wide maps of unselected LOH and CNA tracts in clones derived from S288c x YJM789 hybrid diploids. **A** Experimental design: A total of 30 YPD cultures (15 of the Chr4 plus Chr5 strain JAY2357, and 15 of the Chr13 plus Chr5 strain JAY2358) were started from single cells, grown into colonies, and then transferred to 5 ml tubes until saturation (∼3×10^8^ cells per culture; ∼29 cell generations). Four appropriate aliquot dilutions from each culture were plated, one each: non-selectively to YPD, selectively for single LOH on 5-FOA, selectively for single LOH on canavanine, and selectively for double LOH on 5-FOA plus canavanine. One colony per plate was isolated and whole genome sequenced, for a total of 30 sets of 4 culture-matched clones. **B** shows the results of the WGS analysis. The top horizontal line is the linear end-to-end depiction of the 16 chromosomes in the S288c x YJM789 hybrid diploid strain background, with HetSNPs represented as red S288c alleles and blue YJM789 alleles. Chromosome numbers are indicated above, and black circles represent their respective centromere positions. Each horizontal line corresponds to the genomes of clones that displayed at least one unselected LOH or CNA tract, grouped according to their selection category (no selection; single selection on 5-FOA or Can; double selection on 5-FOA plus Can). None of the no selection control clones contained any LOH or CNA tracts, thus none of those clones are displayed. For the single and double selection groups, the number of unselected tracts in each clone is shown between parentheses. Identical LOH tracts that were detected in all four clones in a matched set (pre-culture) are displayed in the plots, but were not counted toward the total of unselected tracts that arose during each culture. Markers that remained heterozygous were omitted to emphasize visualization of tracts. HetSNPs that were homozygous YJM789/YJM789 (blue) or S288c/S288c (red) are shown as double vertical lines above and below the black chromosome line. No segmental CNAs were found, but multiple whole chromosome CNAs were. Chromosome losses (monosomy) are shown in yellow, chromosome gains (trisomy) are shown in green. Cases of copy-neutral LOH spanning whole chromosomes (uniparental disomy) are shown in orange. Details of all selected and unselected tracts are also available in File S2, Fig. S4-S6, and Table S5. Plots were generated to scale as in Fig. 1B. For size reference, Chr1 is 230 kb.

The quantitative analyses above consistently indicated a degree of coincidence between LOH events at different regions of the genome caused by SGI, however, our estimation of the magnitude of this process (*i.e.*, the excess ratio of observed to expected double LOH rates) was reliant on our ability to accurately measure the single LOH rates used to calculate the expected double rate. Several factors are known to influence the accuracy of mutation rate measurements (29–31), including the fraction of the culture that is plated onto selective media and the method that is used in the calculation itself. Because the rates of single vs. double LOH differed by 2-3 orders of magnitude (Fig. 3C-E), we needed to plate highly diluted cultures (∼100 fold) in order to obtain countable numbers of isolated colonies on single selection Petri dishes, while needing to plate much higher fractions of the same culture onto double selection in order to find a few double LOH colonies. Another possible limitation was that the L&C rate calculation method (23) is susceptible to distortion when the number of mutation events in the culture is too high (*i.e.*, high rate single LOH events in larger than necessary cultures), whereas the Ma-Sandri-Sarkar maximum likelihood estimation method (MSS-MLE; (32) is robust across a wide rate range. Therefore, in order to improve the accuracy and strengthen the rigor of our analysis, we conducted a new round of experimental single LOH rate measurements by using extremely small cultures (20 μl; Methods) grown in microwell plates and plated nearly entirely to selective media, followed by rate calculations using both L&C and MSS-MLE. These procedural refinements resulted in only minor differences in the calculated single LOH rates (Fig. S3 and Table S4). The reduction in the size of cultures resulted in single LOH rate estimates that were equal or at most ∼2 fold higher, and calculations using MSS-MLE produced single LOH rates equal or at most ∼3 fold higher than L&C. In all cases, across all strains, culture volumes and calculation methods, the observed rate of double LOH was always higher than expected, within an excess range of 14 up to 150 fold.

### Characterization of systemic genomic instability in densely heterozygous diploids

We next explored the densely marked genome of the hybrid strain background to carry out a comprehensive characterization of coincident unselected structural variation in clones recovered from the culture-matched LOH rate assays described above. A total of 120 clones were derived from 30 independent cultures, 15 from each hybrid parent (Fig. 3A-B and Fig. 4A). From each culture, we recovered and sequenced a matched set of four clones: one control clone from YPD permissive plates (no selection), one 5-FOA resistant single LOH selection on Chr4 or Chr13, one canavanine resistant single LOH selection on Chr5, and one 5-FOA and canavanine resistant clone from double LOH selection.

This experimental design allowed us to control for any tracts that were present in the precursor cell that gave rise to each culture, before the start of each experiment. Indeed, regions of pre-existing LOH were identified in seven cultures (Table S5, File S2). These identical tracts were present in all four matched sequenced clones from these cultures. Since these tracts were pre-existing and did not accumulate during the growth of the cultures, they were not counted toward the totals described below. In addition, this set-up allowed us to infer the sequential temporal formation relationship between tracts in cases where identical tracts were found in more than one clone in a culture-matched set (see below). A final important feature of this experimental design was that all sequenced clones, regardless of the no, single or double selection regimens, underwent a similar number of cell generations (∼29) since the culture’s precursor cell. This was approximately half the number of generations required earlier to isolate the JAY270-derived smooth and rough clones (Fig. 2), thus restricting the window of opportunity for the sequential accumulation of tracts.

Genomic analysis of the control clones isolated from permissive YPD plates showed that the baseline rates of unselected LOH and CNA were very low under the growth conditions utilized. No unselected tracts were identified in any of the 30 control clones, whereas unselected tracts were detected in several of the clones derived from single and double selection. Excluding the selected tracts on Chr4, Chr13 or Chr5, our analysis revealed the presence of 7 unselected tracts in 5 of the 60 clones (8.3%) derived from single selection, and 14 unselected tracts in 9 of the 30 clones (30%) derived from double selection (Fig. 4B). These results indicated that the likelihood of identifying clones carrying unselected tracts was dependent on whether or not selection for LOH elsewhere in the genome was applied to isolate the clones (p=0.020 from Fisher exact test). Further, double selection led to a stronger enrichment for clones carrying unselected tracts than single selection (p=0.012), consistently with a more robust manifestation of the SGI phenomenon among the double selection clones.

For 3 of the 30 culture-matched clone sets we were able to infer a sequential temporal formation relationship between the selected tracts by examining their endpoint positions (cultures 5, 15 and 27 in Table S5; File S2). For example, in culture 15, the Chr4 LOH tracts were identical in the clones derived from single Chr4 LOH (clone NS30) and double Chr4 plus Chr5 LOH (clone NS60) selections. This suggested that the shared Chr4 LOH event must have occurred first in a common ancestor, followed by the Chr5 loss (monosomy) event detected in NS60. We also applied this same genealogy rationale to analyze the unselected tracts, but found that all were private to the clones where they were detected (none were shared between any of the sequenced matched clones). This was consistent with most unselected tracts possibly appearing in the lineage either at the same time (*i.e.*, SGI burst) or after the formation of the selected tracts present in those respective clones.

The unselected LOH tracts detected in the hybrid diploid background were qualitatively similar to those found in the JAY270 smooth and rough clone sets, with approximately the same 1:1 ratio of interstitial to terminal tracts, and comparable tract sizes. Specifically, there were nine interstitial GC-type LOH tracts (median size 5.0 kb) and seven terminal crossover-type LOH tracts (median size 89 kb). In addition to segmental LOH, we also found cases of unselected genomic alterations spanning whole chromosomes, including monosomy (Chr10) and trisomies (Chr4, Chr10, Chr11, Chr16). Interestingly, and consistent with our recent report (17), four of the five unselected aneuploidies coincided with selected Chr5 monosomy or uniparental disomy (UPD; see clone NS101; Fig 4B), suggesting the possibility of a transient and systemic perturbation of a discrete genome stability pathway (*e.g.*, spindle assembly checkpoint) being responsible for both the selected and unselected aneuploidies detected in these clones.

Finally, this dataset provided additional information about the patterns of genomic alterations for selected LOH, specifically in Chr4, Chr13 and Chr5 (tract maps shown in Figs. S4, S5 and S6, respectively). The patterns for Chr4 and Chr13 were similar to that found earlier for Chr12 in JAY270 (Fig. S2). All 30 Chr4 LOH tracts and 27 of 30 Chr13 LOH tracts were terminal, with endpoints somewhere between the centromere and the CORE2 insertion sites. Several of these had shorter tracts of bidirectional LOH near the endpoint, consistent with GC associated with crossing over (20). One particularly interesting and fortuitous such case was that of clone NS49 (Fig. S4) where the CORE2 marker was within the short GC LOH tract (YJM789 homozygosity), whereas the much longer terminal crossover LOH tract was in the opposite direction (S288c homozygosity) compared to all other selected Chr4 tracts. In addition, three cases of interstitial LOH were found in the Chr13 strain (Fig. S5), with median tract size of 5.4 kb.

The qualitative spectrum of the 60 selected tracts for Chr5 (Fig. S6) was more diverse as it included allelic interhomolog recombination LOH tracts, segmental CNA, as well as whole-chromosome alteration events. The majority (34 of 39) of the allelic recombination-type LOH tracts were terminal, and the remaining (5 of 39) were interstitial with relatively large median tract size of 35.9 kb. One clone (NS59) had a terminal CNA deletion of the left arm of Chr5 spanning *CAN1* with an endpoint at a Ty repeat, coupled with a terminal CNA amplification of the right arm of Chr1 also with another Ty at the endpoint. In addition, this clone had a short, bidirectional, copy neutral LOH tract on the immediate proximal side of the Chr5 deletion endpoint, and PFGE (data not shown) showed a rearranged chromosome of size consistent with a Ty-mediated Chr5/Chr1 non-reciprocal translocation. This interesting combination of proximal allelic and distal non-allelic recombination in the same chromosomal rearrangement event suggested a complex mechanism, perhaps initiated as a short allelic interhomolog break-induced replication tract followed by template switching to the non-allelic Chr1 Ty template (33). Finally, we also found 20 cases of whole-chromosome Chr5 LOH, including 15 cases of simple monosomy where the clones lost the *CAN1*-containing YJM789 homolog, and 5 cases of UPD where the clones carried two full copies of the S288c homolog. Notably, this whole-Chr5 selected LOH class was more prevalent (18 of 20) in the double selection clones (Chr4 plus Chr5, and Chr13 plus Chr5) compared to the single selection clones (2 of 20; Chr5 selection only).

Altogether, the results from our comprehensive sets of experiments provide compelling support for the existence of sporadic and transient bursts of mitotic SGI leading to the rapid accumulation of multiple tracts of structural variation in *S. cerevisiae* cells, including some clones with remarkably rearranged genomes, such as JAY664.

## DISCUSSION

### Precedents of coincident recombination and mitotic origin of SGI

This study allowed us to uncover the SGI phenomenon through which multiple LOH events can accumulate rapidly in a cell lineage. We determined using WGS that yeast clones carrying a primary selected LOH tract at any of four different genomic regions were more likely than expected to carry unselected LOH tracts elsewhere. We also showed in quantitative LOH assays that combinations of double LOH at Chr5 and Chr4 or Chr13, occurred at rates 14-150 fold higher than expected if single LOH events always occurred independently. We interpret these results as evidence for the occurrence of bursts of genomic instability leading to multiple LOH events over one or few mitotic cell generations.

Spontaneous mitotic recombination events like the ones described here are triggered by local DNA lesions and/or replication fork collapse episodes, which then lead to chromosomal breakage and HR repair using the allelic homolog or a non-allelic homologous sequence as template (20). Such precursor lesions are thought to occur mostly randomly in vegetative cells, both spatially across the genome and temporally, therefore mitotic recombination events involving different chromosomes should rarely be coincident. In contrast, meiotic recombination is known to be a systemic genetic variation process, since it occurs simultaneously throughout the genome and involves intricate coordination between generation and repair of genome-wide double strand breaks (25).

While our study is, to our knowledge, one of the first to describe the SGI phenomenon through the lens of high-resolution genome-wide analytical methods, there have been sparse reports of elevated coincident mitotic recombination in yeasts as well as in mammalian cells dating back decades (8-10, 13-15, 34, 35). The typical experimental design in those cases was to select clones for carrying a recombination event at a primary locus, and then screening the resulting clones for the occurrence of secondary unselected recombination at one or a limited number of unlinked marked loci. The same intriguing observation, shared in all cases, was a frequency of coincident recombination that was higher than that predicted assuming the individual events always occurred independently.

In some of the yeast studies, the high coincident recombination rates were interpreted as being derived from a small number of cells within the replicating population that spuriously entered the meiotic developmental program, or transiently experienced a “para-meiotic” state, but reverted back to mitotic growth (9, 10). A recent study specifically characterized this type of return-to-growth (RTG) event and the genome-wide recombination outcomes associated with it (24). The authors often detected a large number of LOH tracts per clone (minimum of 5, average of ∼30, and up to 87), indicating that RTG induction leads to abundant and widespread recombination. Another notable finding was that while interstitial GC LOH tracts were frequent, their sizes were relatively constrained (2.3 kb on average). This measurement is notable because it is consistent with GC tract sizes measured in haploids derived from complete meiotic divisions; ∼2 kb median size (36). In contrast, GC tracts associated with mitotic recombination tend to be significantly longer, approximately 5-6 kb median size (37). This increase in typical GC tract sizes is likely a reflection of subtle mechanistic differences in the processing of HR intermediates between meiotic and mitotic cells.

The studies outlined above suggest that cryptic initiation of meiosis in a small number of cells can in some cases lead to coincident recombination, however, we favor the interpretation that the events analyzed in our study originated primarily from *bona fide* mitotic cells. The work by Laureau *et al*. (24) described above clearly defined the features of systemic LOH caused by meiotic initiation followed by RTG. The pattern we detected in our study was different, and instead was consistent with mitotic patterns. The number of unselected interstitial GC LOH tracts per clone we detected was small (typically 1 to 3) and their sizes were long (∼5.0 kb). This was reinforced by the observation that *MAT*a/*matΔ* diploids, incapable of entering the meiotic developmental program, continued to display double LOH rates that were higher than expected from independent events (Fig. 3D).

Our interpretation of a mitotic origin for SGI is supported by other reports of high coincident recombination in proliferating cells in which the induction of a full meiotic cycle, RTG or para-meiosis were either unlikely or could be ruled out entirely. One study in *S. cerevisiae* specifically measured the formation of spurious haploids from mitotic diploid cultures displaying high coincident intragenic recombination at unlinked pairs of heteroalleles (14). The authors found that while haploids did form in their cultures, the frequency was far below that needed to influence the formation of double recombinants, thus concluding that a low level of cryptic meiosis was not a likely contributor. In addition, one of the seminal studies (15) of LOH in *C. albicans* (a species devoid of a conventional sexual cycle (38)), reported data that closely parallel our own observations. First, the authors selected clones for the presence of a primary LOH event at the *GAL1* locus on chromosome 1. Then, using a SNP-array platform, they detected frequent unselected secondary LOH tracts among clones carrying the primary event, but rarely in control clones still heterozygous at *GAL1*. In addition, selection for LOH at *GAL1* was associated with the emergence of altered colony morphology phenotypes, presumably derived from rearrangements elsewhere in the genome. Accordingly, clones displaying altered morphology were enriched for the presence of unselected LOH tracts when compared to clones with normal morphology. A recent expanded study by the same group corroborated their original findings using high resolution sequencing-based approaches (16).

Another important pair of precedents of mitotic SGI observations comes from experiments conducted in mammalian systems. These used either human TK6 lymphoblastoid cells in culture (34), or mouse kidney cells *in vivo* and in culture (35). In both cases, the starting cells were heterozygous for mutations at the counter-selectable markers, *TK* and *Aprt*, respectively, enabling the selection of clones carrying a primary LOH event at those loci. Subsequently, the presence of secondary LOH tracts was assessed at roughly a dozen marker loci elsewhere in the human or mouse genomes. The two studies found that secondary LOH was more frequent in clones selected for carrying the primary LOH event than in controls clones that remained heterozygous. These studies demonstrated that SGI also likely exists in metazoans, and can be detected in cells that are exclusively mitotic.

### SGI-like observations in human disease

In addition to the experimental examples above, our results also resemble recent reports of bursts of mitotic genomic instability in humans during cancer genome evolution and early development. Specifically, genome-wide copy number profiling of thousands of individual cells isolated from tumors from patients with triple-negative breast cancer revealed that a large number of CNAs were acquired within a short period of time at the early stages of tumor development (3). Most of these CNAs were shared between several cells from a same tumor, suggesting the occurrence of a burst of genomic instability in one or few initiating cells followed by a long period of stable clonal expansion. Although the study had power to detect gradual accumulation of mutations, no clones with intermediate CNA profiles were identified, suggesting a punctuated model of mutation accumulation (1, 2). This same conclusion has been corroborated recently through spatially resolved breast tumor single cell sequencing (39), in colorectal cancer (4), and even more broadly in thousands of tumor samples from dozens of different cancer types reported by the pan-cancer analysis of whole genomes (PCAWG) consortium (6).

Another pertinent parallel is the recent analysis of patients with genomic disorders that carry multiple *de novo* constitutional CNVs (MdnCNVs; (7)). Typically in those patients, only one of the structural variants was the primary event causing the symptoms associated with the disorder. The additional CNVs were secondary, occurred at unrelated regions, and apparently formed during a short burst of genomic instability at some point in the perizygotic time interval. The changes then propagated stably during development to be found in all cells in the patients. Taken together, these results suggest that SGI processes may be universal and can play an important role in human disease development.

### Possible mechanisms underlying SGI

While our results support a pronounced contribution of SGI to the rapid accumulation of structural variation in budding yeast, the specific causes for the existence of a small subset of recombination-prone cells within a normal mitotic population remain to be determined. This phenomenon most likely has multiple and distinct origins, however, we favor two non-exclusive mechanisms, related to cellular ageing and stochastic gene expression. These two models are attractive because they are transient in nature, which would support stable transmission of rearranged genomic structures after the systemic vulnerability time window has passed.

The first scenario is that clones carrying multiple unselected LOH events originated from replicatively old mother cells. This model stems from the observation of a marked increase in the rate of LOH in daughter yeast cells budded from mothers that had undergone ∼25 cell divisions (40), relatively old within the context of a typical maximum *S. cerevisiae* replicative lifespan of ∼38-40. Subsequent work showed that this increase in nuclear genomic instability was strongly correlated with the initial appearance of mitochondrial DNA loss and/or damage in the old mother cells (41). In our study, however, all of the spontaneous rough colony isolates analyzed retained normal respiratory activity (all were non-*petite*; grew on non-fermentable carbon sources), so they must have had integral mitochondrial genomes. They also did not show signs of continual genomic instability. Therefore, if replicative aging were an underlying factor in SGI, it would be through a pathway that does not involve loss of mitochondrial function.

Another explanation for a subpopulation of transiently hyper-recombinogenic cells involves heterogeneities that exist within an isogenic population. Specifically, cell-to-cell variation (*i.e.* noise) in gene expression has been reported in organisms ranging from prokaryotes, to yeast, to humans (42). It is plausible that stochastic variation in the expression of a broad class of genes involved in genome stability could cause specific protein levels to drop below or rise above those required for optimal function. A recent comprehensive genome stability network analysis identified 182 genes involved in suppression of gross chromosomal rearrangements (43), and an earlier genetic screen identified 61 genes specifically involved in suppressing LOH (44). In this scenario, rare individual cells that fail to adequately express any of these genes could effectively behave as defective mutants and display a mutator phenotype for a short period. Some of these genes act cooperatively, therefore concomitant loss of activity causes extreme levels of genomic instability. For example, double knockouts for *TEL1* and *MEC1*, encoding critically important DNA damage response proteins (orthologs of mammalian ATM and ATR, respectively), show marked increase in mitotic genomic instability (45), often accumulating multiple genome rearrangements (46). A similar extreme phenotype might be expected in a wild type cell that by chance simultaneously had a critically low level of transcription of these two genes.

Likewise, overexpression of single genes encoding a subunit of a genome stability multi-protein complex, or a critical regulatory component of a DNA repair reaction, could also lead to a dominant negative phenotype that temporarily impairs function. A recent study (47) demonstrated precisely this effect in budding yeast by sorting wild type cells that randomly over or under-expressed *RAD52* or *RAD27*. The individual cells at the extremes of the expression spectrum had markedly altered rates of recombination. Importantly, the hyper-recombinogenic state of these individuals would be completely reversible once the descendant cells returned to the gene expression levels typical of most individuals in the population. Another way through which noise in gene expression could lead to transient increases in structural variation and all other forms of mutation, including base substitutions (48), would be by overexpression of genes involved in the formation of endogenous DNA damage (49). In this case, cells would be overburdened by systemic genome-wide damage for a brief period leading to the simultaneous accumulation of multiple seemingly independent mutations. This mechanism could explain the observed stable clonal expansion that followed SGI in our selected LOH clones, as well as in the recent *in vivo* human studies above.

The genomic analyses of clones carrying primary selected LOH tracts described in our study provides detailed information about the nature and frequency of secondary recombination events resulting from the SGI process. Our study also provides a unifying context for the interpretation of classic and recent reports of coincident recombination in yeasts, in mammalian experimental systems, and in human disease. The combination of whole genome analyses, modern lineage tracing and single cell transcriptomic approaches (50, 51), and the double LOH selection approach described here offer a powerful experimental platform to further dissect the core mechanisms responsible for the SGI phenomenon.

## MATERIALS AND METHODS

### Yeast genetic backgrounds, growth media and procedures

*Saccharomyces cerevisiae* strains used in this study descended from the JAY270 (52) or CG379 (53) strain backgrounds, or from crossings between the strains S288c and YJM789 (Table S1). Standard procedures for yeast transformation, crossing and sporulation were followed (54). Cells were grown in YPD and synthetic minimal media (SC) at 30C, under rotation for liquid cultures. *URA3, KanMX4, NatMX4* and *HphMX4* transformants were selected in uracil drop-out SC, YPD plus 400 mg/L Geneticin, 200 mg/L of Nourseothricin (Nat) and 300 mg/L of Hygromycin B, respectively. Counter-selection against *URA3* and *CAN1* were performed in SC plus 1 g/L of 5-FOA and 60 ml/L of canavanine in arginine drop-out, respectively.

### Isolation of smooth and rough clones derived from JAY270

The control smooth clones were isolated by inoculating 28 independent 2-day old single smooth colonies from YDP agar plates onto 5 ml of YPD liquid. Liquid cultures were grown until saturation for 24 hours at which point 50 μl (∼5×10^6^ cells) were transferred to fresh 5 ml YPD to start a new passage cycle. At the end of the fifth consecutive liquid passage the cultures were diluted, plated onto YPD and a single random smooth colony was isolated and frozen for genomic analysis. We estimated this growth regimen involved ∼57 consecutive cell generations (22 in agar, plus 35 in liquid [5 passages x 7 generations per passage]). The 23 independent rough colonies isolated in this study were derived through a similar growth protocol, but aliquots were plated to YPD for visual screening of colony morphology at the end of every liquid growth cycle. The liquid passages were discontinued once the first rough colony was identified and frozen from each culture. The estimated median number of cell generations after which rough clones were isolated was ∼50 (Table S3). For the five rough clones isolated previously (18), including JAY664, and three of the rough clones isolated in this study, the total number of liquid passages was not recorded. However, because we never exceeded ten passages, the number of cell generations associated with these clones was at most ∼92 (22+[10×7]), likely less.

### Strain construction for LOH assays

The JAY270 strains used in the LOH assays were constructed from a homozygous *ura3/ura3* derivative of JAY270 (JAY585; gift from F. Galzerani). The CORE2 cassette containing the *Kluveromyces lactis URA3* gene, the *S. cerevisiae URA3* gene and the *KanMX4* geneticin resistance marker (*KlURA3-ScURA3-KanMX4*) was amplified from pJA40 (55) with primers targeting two genomic regions (Table S2). Primers JAO506 and JAO507 were used for integration of CORE2 distal to *SSF2* in Chr4, and JAO502 and JAO503 for integration proximal to *ADH6* in Chr13. Transformation of JAY585 with each cassette resulted in JAY865 (*SSF2*/*SSF2*::CORE2) and JAY868 (*ADH6*/*ADH6*::CORE2), which were used in the single 5-FOA^R^ LOH assays. Subsequently, one native copy of the *CAN1* gene was deleted from each of these strains using the *NatMX4* cassette. The cassette was amplified from pAG25 (56) using primers JAO271 and JAO272 and transformed into JAY865 and JAY868, resulting in JAY1804 and JAY1812, respectively. These strains were used for Can^R^ single LOH assays and for Can^R^ - 5-FOA^R^ double LOH assays. The same procedure was followed to build LOH assay strains in the CG379 background, resulting in strains JAY861 and JAY859 (5-FOA^R^ single LOH assays), and strains JAY1567 and JAY1569 (Can^R^ single LOH assays and Can^R^ - 5-FOA^R^ double LOH assays). We further manipulated JAY1567 and JAY1569 to create *MATa/matΔ* isogenic derivatives. The *HphMX4* cassette was amplified from pAG32 (56) using the primers JAO1440 and JAO1441. This cassette was used to replace a segment of the *MATα* allele in JAY1567 and JAY1569, resulting in JAY1808 and JAY1809 respectively. A similar procedure was used to build hybrid strains in the S288c/YJM789 background used in the LOH assays. In this case, JAY297 (S288c *MATa*) (57) was transformed for integration of the CORE2 cassette distal to *SSF2* in Chr4 and proximal to *ADH6* in Chr13. Subsequently, the native copy of *CAN1* was replaced with a *NatMX4* cassette in each strain, resulting in JAY2355 and JAY2356, respectively. These strains were then crossed to JAY308 (YJM789, *MATα),* resulting in JAY2357 (*MATa/MATα, SSF2*/*SSF2*::CORE2 *CAN1/can1Δ*) and JAY2358 (*MATa/MATα, ADH6*/*ADH6*::CORE2 *CAN1/can1Δ*).

### Quantitative LOH rate assays

For conventional culture size recombination assays, single colonies were inoculated into 5 ml liquid YPD, and incubated for 24 hours at 30C in a rotating drum. For small culture LOH assays, 5 ml cultures were inoculated and incubated at 30C for 24 hours. Cultures were then diluted in fresh YPD to a final concentration of ∼30 cells/ml, and 20 μL aliquots of this dilution were dispensed into individual wells of 96 well plates and incubated at 30C for 48 hours. At this concentration, most wells did not show any growth, whereas wells with growth were likely started from a single cell. The cultures were serially diluted and plated on YPD (permissive), and SC plus 5-FOA (selective) and/or canavanine (selective). Colony counts were used to calculate recombination rates and 95% confidence intervals using the Lea & Coulson method of the median within the FALCOR web application (23, 58). Recombination rates and 95% confidence intervals were also calculated through the Ma-Sandri-Sarkar maximum likelihood estimation method (MSS-MLE) with plating fraction correction using the FluCalc web application (31).

### Genome Sequencing Analyses

The Illumina short read WGS platform was used to sequence the genomes of the JAY270 derivatives (28 smooth and 28 rough clones) and 120 derivatives from theS288c/YJM789 hybrid strain. Oxford Nanopore single molecule long reads were also generated for JAY270 and JAY664. All genome sequencing data associated with this study is available in the Sequence Read Archive (SRA) database under study number SRP082524.

#### Detection of LOH tracts in the JAY270 background

LOH tracts were detected using CLC genomics workbench software to map sequencing reads from the smooth and rough clones onto the S288c reference and detecting SNPs across the whole genome. Low stringency detection parameters were set such that SNPs present at frequencies higher than 0.05 were identified. We then interrogated the 12,023 loci in the JAY270 HetSNP list generated previously (18), determining the nucleotides present at those positions and their relative frequency. When no SNPs were detected at those specific positions, the genotype was called as homozygous for the reference nucleotide. When the alternative nucleotide was detected at frequency higher than 0.95 the genotype was called homozygous for the alternative nucleotide. Alternative nucleotides detected at frequencies between 0.1 and 0.9 resulted in a heterozygous call for that locus. After the genotypes were called they were then converted to the respective haplotype designations as M/M and P/P homozygous, and M/P heterozygous. In control analysis using WGS reads from JAY270, all 12,023 loci were called as heterozygous. LOH tract sizes were estimated by calculating the positions of breakpoints to the right and to the left, and subtracting the left side position from the right side. Breakpoint positions were calculated as the average position between the two HetSNPs that defined the transition from heterozygosis to homozygosis. For terminal LOH tracts the coordinates of the left or right telomeres were used as the breakpoint positions. LOH tracts were called even if they included homozygosity of a single marker HetSNP. Six such cases were identified, all were interstitial. All were validated through direct visual inspection of the read mapping files. A subset of these single marker LOH calls were independently validated by PCR and Sanger sequencing, or by Nanopore WGS (Fig. S1).

#### Detection of LOH tracts in the S288c/YJM789 hybrid background

A list of heterozygous loci in the S288c/YJM789 diploid background was generated by combining overlapping SNP data from the strains JAY2355, JAY2356, JAY2357, JAY2358 and JAY308, which resulted in 51,053 HetSNPs. This list was then filtered manually to identify markers commonly identified as false positives, resulting in a final list of 50,708 HetSNPs that were interrogated in the S288c/YJM789*-*derived clones isolated from LOH assays. LOH was called when the nucleotide frequency in a specific position was <0.1 or >0.9 and all LOH calls were supported by 10 or more reads. LOH tract sizes were calculated using the same approach described for the LOH analysis in the JAY270 derivatives above.

#### Detection of CNA tracts in JAY270 and S288c/YJM789 derivatives

Reference genome read mapping BAM files from each of the control and experimental clones generated in CLC genomics workbench were imported into Biodiscovery Nexus Copy Number software. CNA analyses in Nexus were conducted based on read depth of coverage using as reference BAM files generated from read mappings of the appropriate JAY270 or S288c/YJM789 parent strains. In clones where regions of segmental or full-chromosome CNA were identified, the coordinates of these calls were cross-referenced to the LOH calls above to obtain a final classification of CNA tracts as segmental deletion or amplification, and in the case of aneuploidies, as monosomy, uniparental disomy or trisomy.

### Statistical Analyses

To compare the number of unselected LOH tracts in JAY270 smooth vs. rough clones we used a test of equal proportions. We also tested the Poison distribution goodness of fit for the number of unselected tracts per clone within smooth and rough clone sets. In the S288cxYJM789 hybrid system, the Fisher Exact Test was used to determine whether the probability of having unselected LOH tracts was dependent on the initial selection of LOH elsewhere in the genome. We first compared the type of selection used (none vs. any) to the number of clones showing a presence or absence of unselected LOH tracts, yielding a p-value of 0.020. Second, we compared the type of selection used (single vs. double) to the number of clones showing a presence or absence of unselected LOH tracts, yielding a p-value of 0.012. All calculations were done in R.

## ACKNOWLEDGMENTS

We thank Tom Petes, Michael McMurray, Dmitry Gordenin, Aaron Mitchell for valuable insights and comments on earlier versions of the manuscript. NMVS received a pre-doctoral fellowship from Brazil’s CAPES (0316/13-0). PC and AVP were supported by a UGC fellowship. KTN was supported by a Wellcome Trust-DBT India Alliance Intermediate fellowship- (IA/I/11/2500268) and IISER-TVM intramural funds. Research reported here was supported by an NIH grant to JLA (R35GM119788), and an NIH grant to LRH (K99GM134193).

---

**Figure S1.**
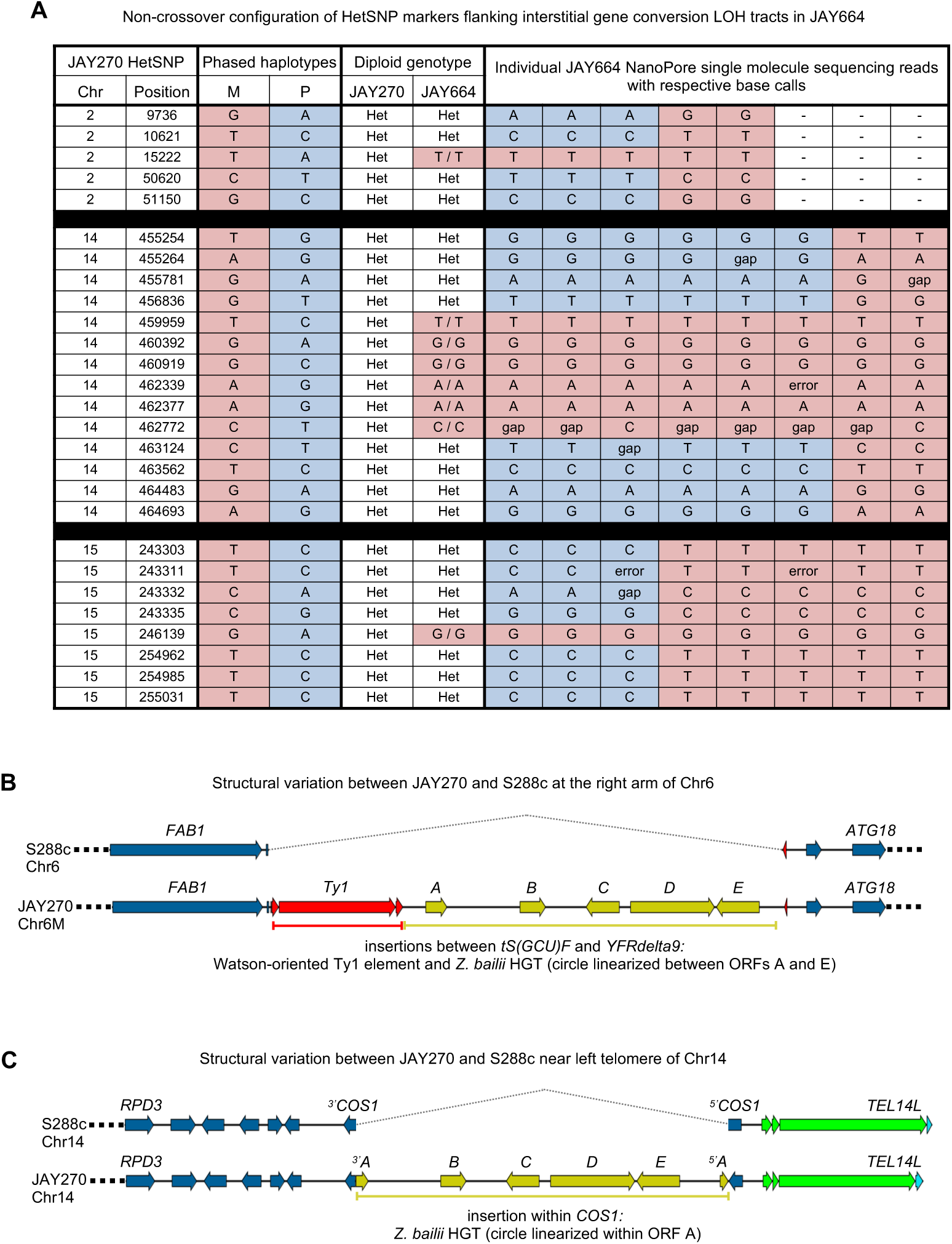
Structural genomic analyses in JAY664 and JAY270 through Nanopore single molecule long read WGS. **A** shows the recombination outcomes associated with the three interstitial LOH tracts detected in JAY664. Each row corresponds to JAY270 HetSNP positions within or flanking the LOH tracts. M alleles are shaded in red, P alleles are shade in blue. LOH tracts in JAY664 are also shaded to highlight the LOH regions (all three were M/M; red). Each column on the right side of the table contains the base calls made for individual single molecule long reads that spanned the entire regions of Chr2 (top), Chr14 (middle), or Chr15 (bottom). For all three LOH tracts, the recombinant homolog (recipient molecule) has P alleles present before and after the (P to M) gene conversion tract, whereas the other homolog (donor molecule) has M alleles present before, within, and after the tract. This pattern demonstrated that the gene conversion event was not associated with exchange of flanking markers (*i.e.*, non-crossover resolution of the recombination intermediate), and ruled out a gene conversion with associated reciprocal crossover mechanism, which would have caused the telomeric-proximal markers to remain heterozygous. **B** and **C** show respectively, the positions and ORF arrangements of the *Zb* circle insertions in JAY270 Chr6-M and Chr14 relative the S288c reference genome. The color pattern follows the system described in the legend of Fig. 1C.

**Figure S2.**
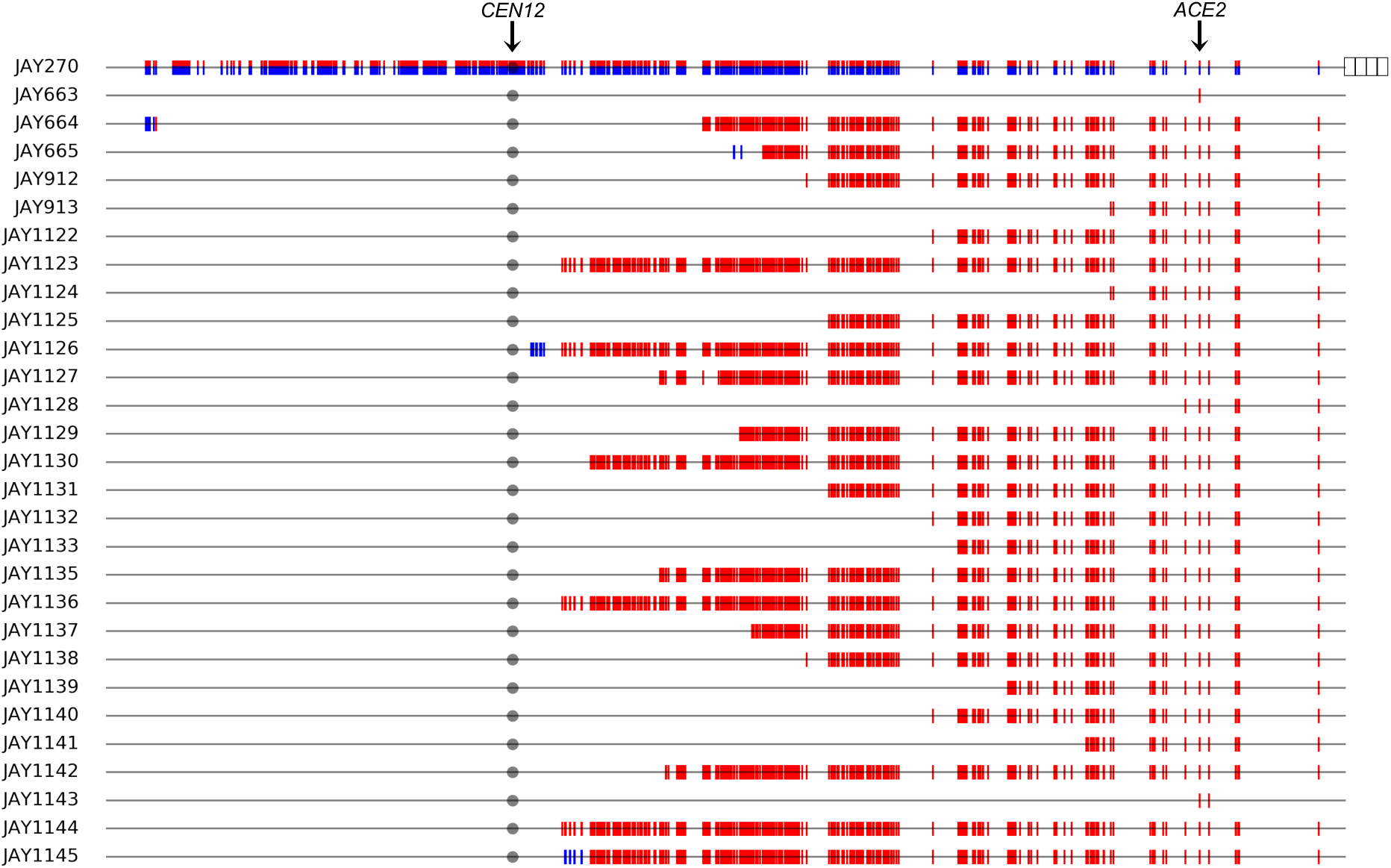
Maps of selected Chr12 LOH tracts spanning the *ACE2* locus in rough colony clones. The top horizontal line is the linear depiction of Chr12 in the JAY270 strain, from the left telomere to the rDNA cluster (right; striped boxes), with HetSNPs represented as paternal (blue) and maternal (red) markers. Note that Chr12 regions distal to the rDNA do not contain any heterozygous markers in JAY270 and thus are not shown. The position of the *ACE2* locus is shown. Each horizontal line below corresponds to the selected Chr12 LOH tract maps in each of the rough clones. HetSNP positions that were homozygous P/P (blue) or M/M (red) are shown as double vertical lines above and below the black chromosome line. Markers that remained heterozygous M/P were omitted to emphasize visualization of LOH tracts. As expected from selection for the rough colony morphology, all rough clones were homozygous for the maternal *ace2-A7* allele (red). Plots were generated to scale in Python 2.7 using the matplotlib package and a custom script. For size reference, the distance between the left telomere and the right-most HetSNP (proximal to the rDNA) is 450 kb. The predominant class (26 of 28) of selected Chr12 LOH tracts spanning the *ACE2* locus in rough colony clones included tracts that were terminal (extended from a proximal position between *CEN12* and *ACE2*, up to the end of Chr12 heterozygosity near the rDNA cluster). Four of these clones had complex discontinuities and/or showed limited reversed LOH pattern for the Chr12-P alleles near the endpoint. Interstitial tracts spanning only a limited region that included the *ACE2* locus were found in only two of the rough clones. These two broad tract classes, terminal and interstitial, were consistent with crossover and gene conversion-only outcomes of interhomolog mitotic recombination, respectively. The large overall excess of crossover-type relative to gene conversion-type recombination outcomes (26:2) was similar to that described in our previous characterization of selected Chr12 LOH tracts in JAY270 (18).

**Figure S3.**
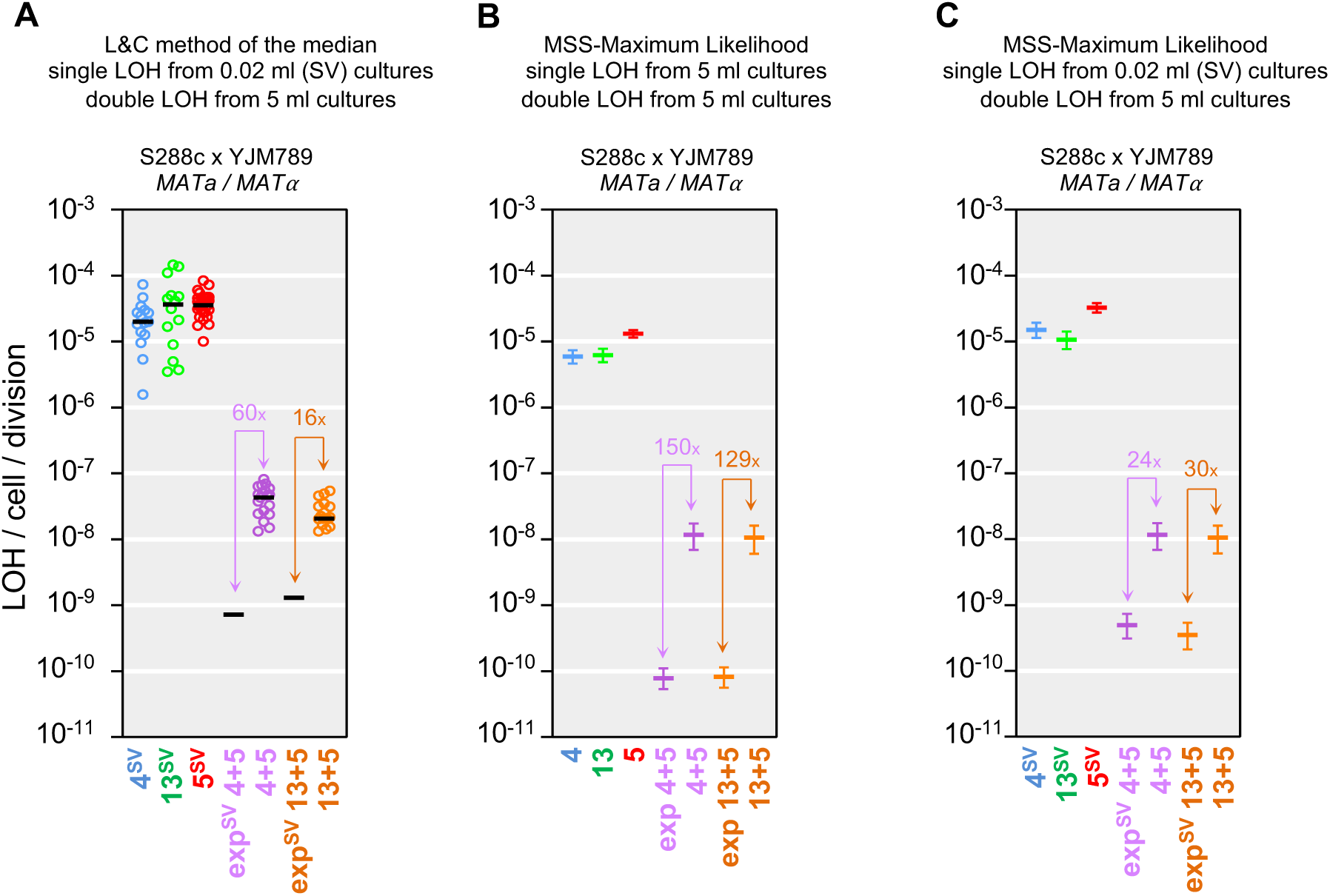
Alternative approaches to the calculation of single and double LOH rates in the S288c x YJM789 hybrid strain background. **A** shows LOH rate data calculated using the Lea & Coulson method of the median. In this case, the single LOH rates for Chr4, Chr13 and Chr5 were obtained from small volume (SV) cultures (0.02 ml YPD), while the double LOH rates were obtained from regular volume cultures (5 ml YPD; same values plotted in Fig. 3E). “exp^SV^” in the Y-axis designates the expected double LOH rates based on the multiplicative product of the pertinent SV median single LOH rate pairs. The excess fold ratio of experimentally observed over calculated expected double LOH rates are shown in all pairwise comparisons. Circles indicate the experimentally determined rates in individual cultures, color-coded according to single chromosomes as in Fig. 3A and 3B, and double chromosomes using purple for Chr4 plus Chr5 or orange for Chr13 plus Chr5. Black horizontal bars indicate the median rate values for each culture set. Only the black bar is shown in the “exp^SV^” cases. The same numerical data are shown in Table S6, including 95% confidence intervals (95% CI; not plotted in **A**). **B** shows single and double LOH rates and 95% CI calculated using the MSS-maximum likelihood (MSS-MLL) method with dilution volume correction from the same colony count data used for Fig. 3E (regular 5 ml YPD cultures for single and double LOH). Note that (unlike Lea & Coulson) the MSS-MLL method does not produce mutation rates for individual cultures, just a rate and 95% CI (plotted following color code above) for the collective set of culture repetitions in each measurement. The expected double LOH rates and 95% CI were derived by multiplying the pertinent 5 ml MSS-MLL single LOH rates. **C** shows single and double mutation rates and 95% CI calculated using the MSS-MLL method with dilution volume correction using SV 0.02 ml YPD for single LOH and regular 5 ml YPD double LOH (same values as in **B**) cultures. The expected SV double LOH rates and 95% CI were derived by multiplying the pertinent SV 0.02 ml MSS-MLL single LOH rates.

**Figure S4.**
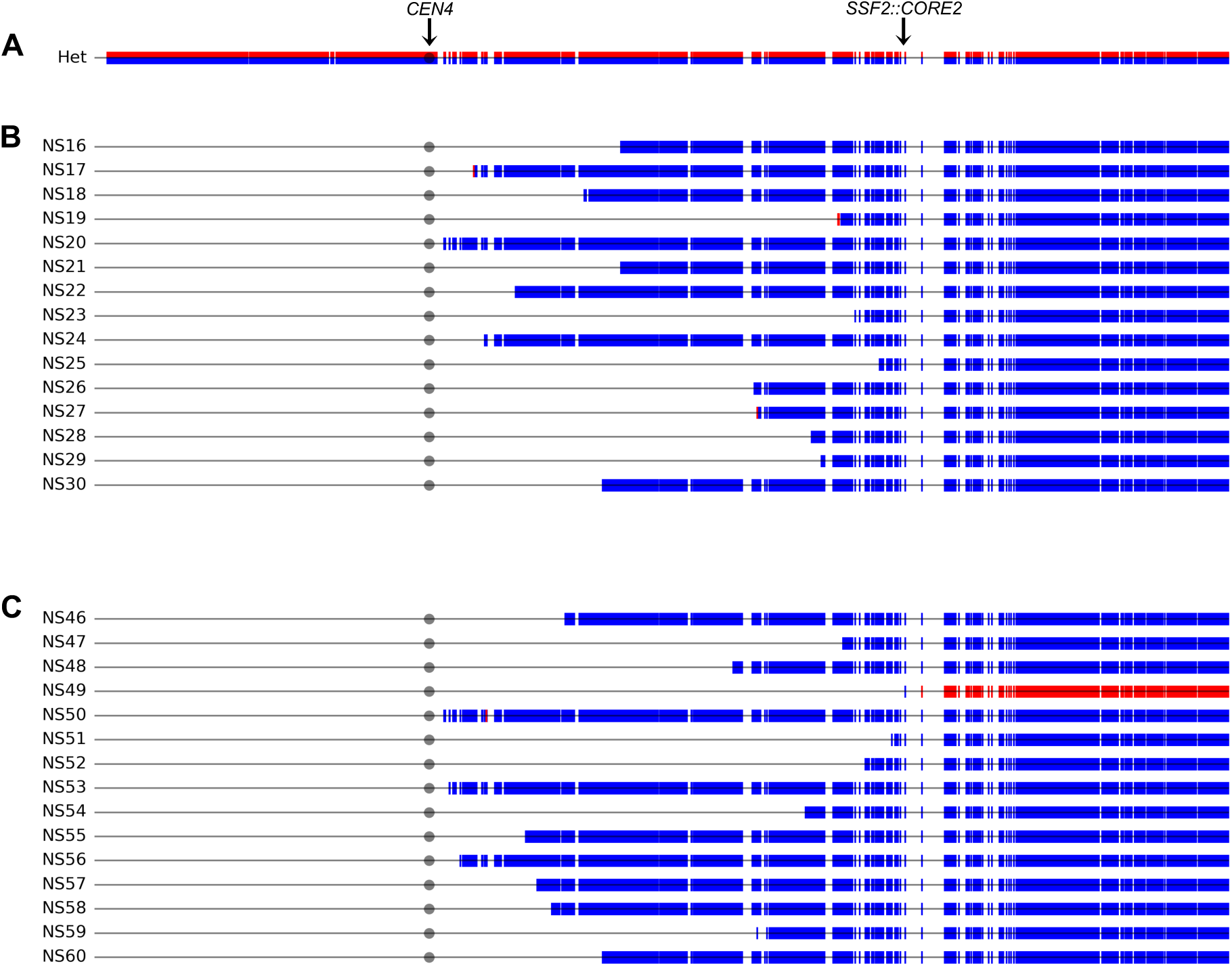
Maps of selected Chr4 LOH tracts spanning the *SSF2::CORE2* insertion locus. The horizontal line shown in **A** is the linear depiction of Chr4 in the S288c/YJM789 hybrid strain background, with HetSNPs represented as red S288c alleles and blue YJM789 alleles. The circle indicates the position of the centromere and the arrow indicates the position of the *SSF2::CORE2* insertion. Each horizontal line in **B** and **C** below corresponds to the selected Chr4 LOH tract maps in each of the single and double selection clones, respectively. Markers that remained heterozygous were omitted to emphasize visualization of tracts. HetSNPs that were homozygous YJM789/YJM789 (blue) or S288c/S288c (red) are shown as double vertical lines above and below the chromosome line. Details of all selected and unselected tracts are also available in File S2 and Table S5. Plots were generated to scale in Python 2.7 using the matplotlib package and a custom script. For size reference, Chr4 is 1525 kb.

**Figure S5.**
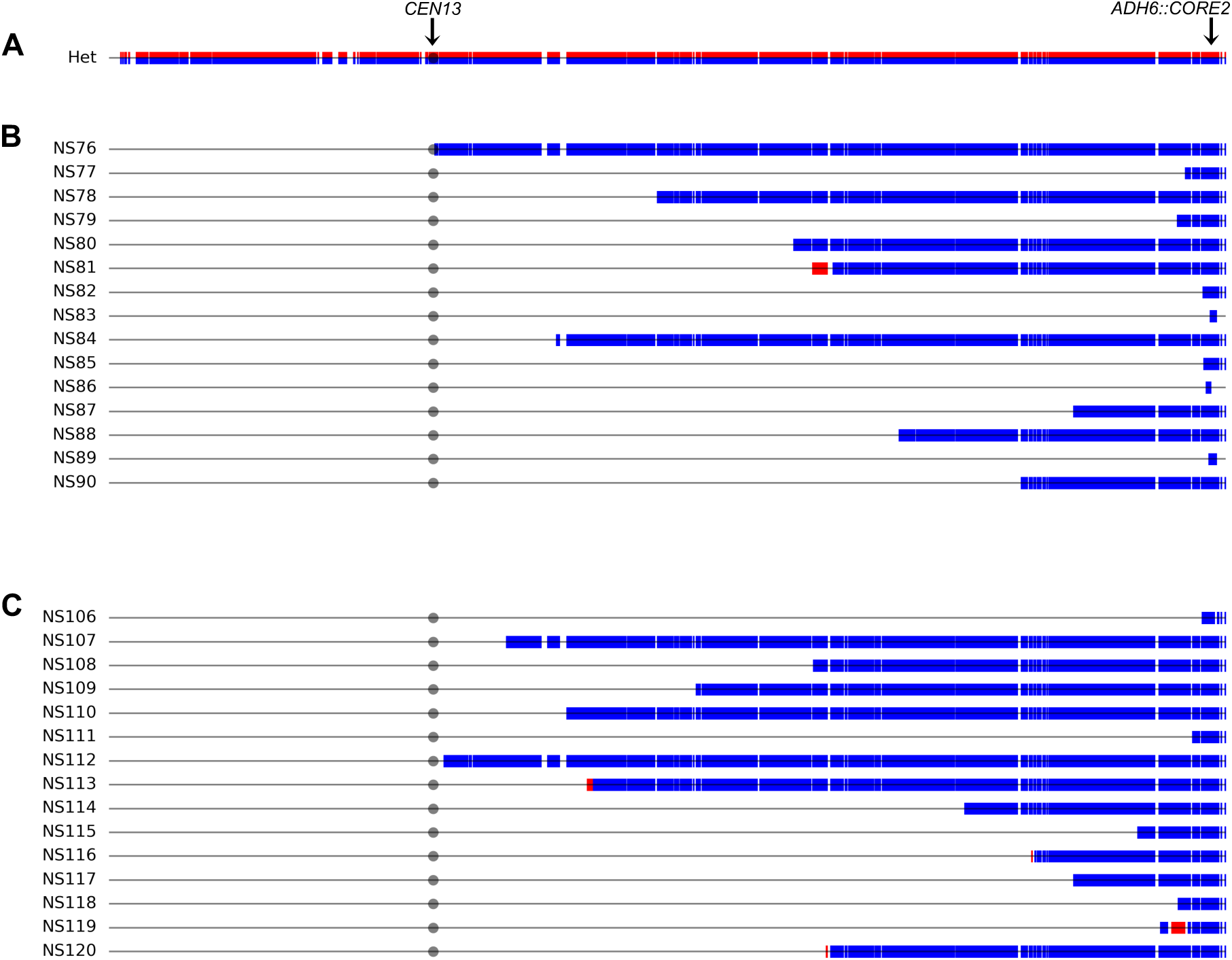
Maps of selected Chr13 LOH tracts spanning the *ADH6::CORE2* insertion locus. The horizontal line shown in **A** is the linear depiction of Chr13 in the S288c/YJM789 hybrid strain background, with HetSNPs represented as red S288c alleles and blue YJM789 alleles. The circle indicates the position of the centromere and the arrow indicates the position of the *ADH6::CORE2* insertion. Each horizontal line in **B** and **C** below corresponds to the selected Chr13 LOH tract maps in each of the single and double selection clones, respectively. Markers that remained heterozygous were omitted to emphasize visualization of tracts. HetSNPs that were homozygous YJM789/YJM789 (blue) or S288c/S288c (red) are shown as double vertical lines above and below the chromosome line. Details of all selected and unselected tracts are also available in File S2 and Table S5. Plots were generated to scale in Python 2.7 using the matplotlib package and a custom script. For size reference, Chr4 is 1525 kb.

**Figure S6.**
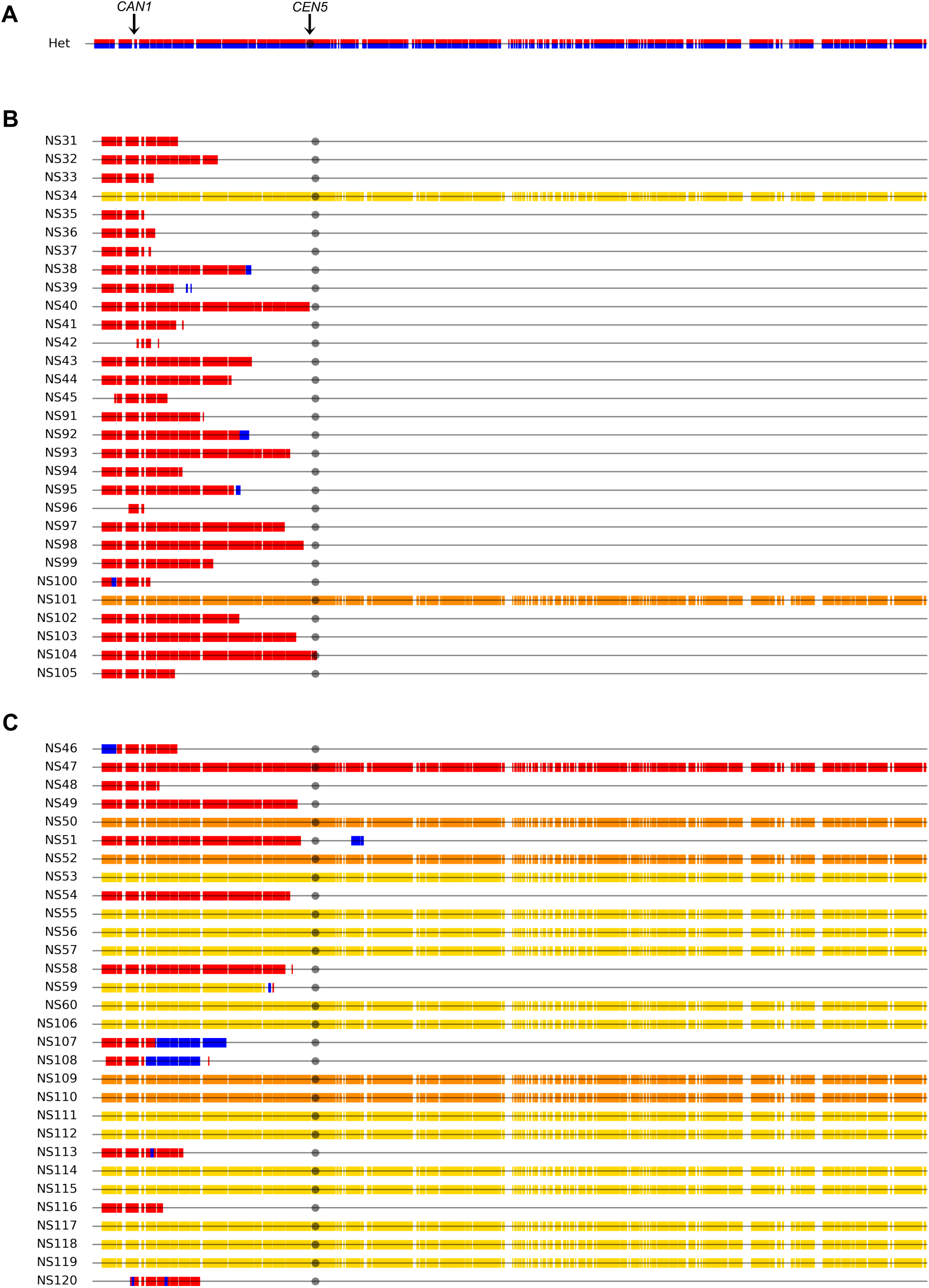
Maps of selected Chr5 LOH tracts spanning the *CAN1* locus. The horizontal line shown in **A** is the linear depiction of Chr5 in the S288c/YJM789 hybrid strain background, with HetSNPs represented as red S288c alleles and blue YJM789 alleles. The circle indicates the position of the centromere and the arrow indicates the position of the *CAN1* locus. Each horizontal line in **B** and **C** below corresponds to the selected Chr5 LOH tract maps in each of the single and double selection clones, respectively. Markers that remained heterozygous were omitted to emphasize visualization of tracts. HetSNPs that were homozygous YJM789/YJM789 (blue) or S288c/S288c (red) are shown as double vertical lines above and below the chromosome line. Whole chromosome alterations are shown in solid colors as follows: Chromosome losses (monosomy) are shown in yellow and cases of copy neutral LOH spanning the Chr5 (uniparental disomy; UPD) are shown in orange. Details of all selected and unselected tracts are also available in File S2 and Table S5. Plots were generated to scale in Python 2.7 using the matplotlib package and a custom script. For size reference, Chr4 is 1525 kb.

**Table S1.**
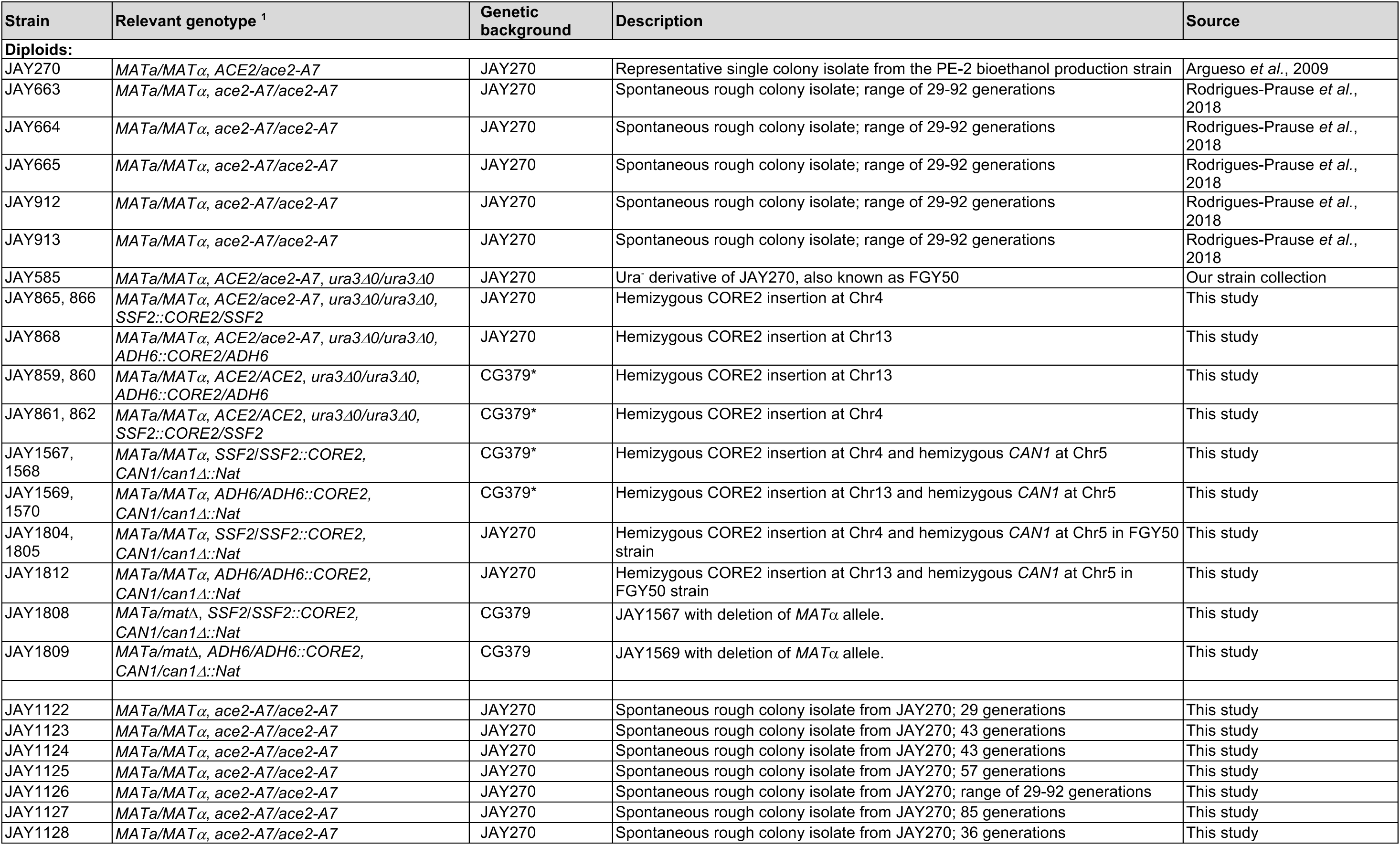
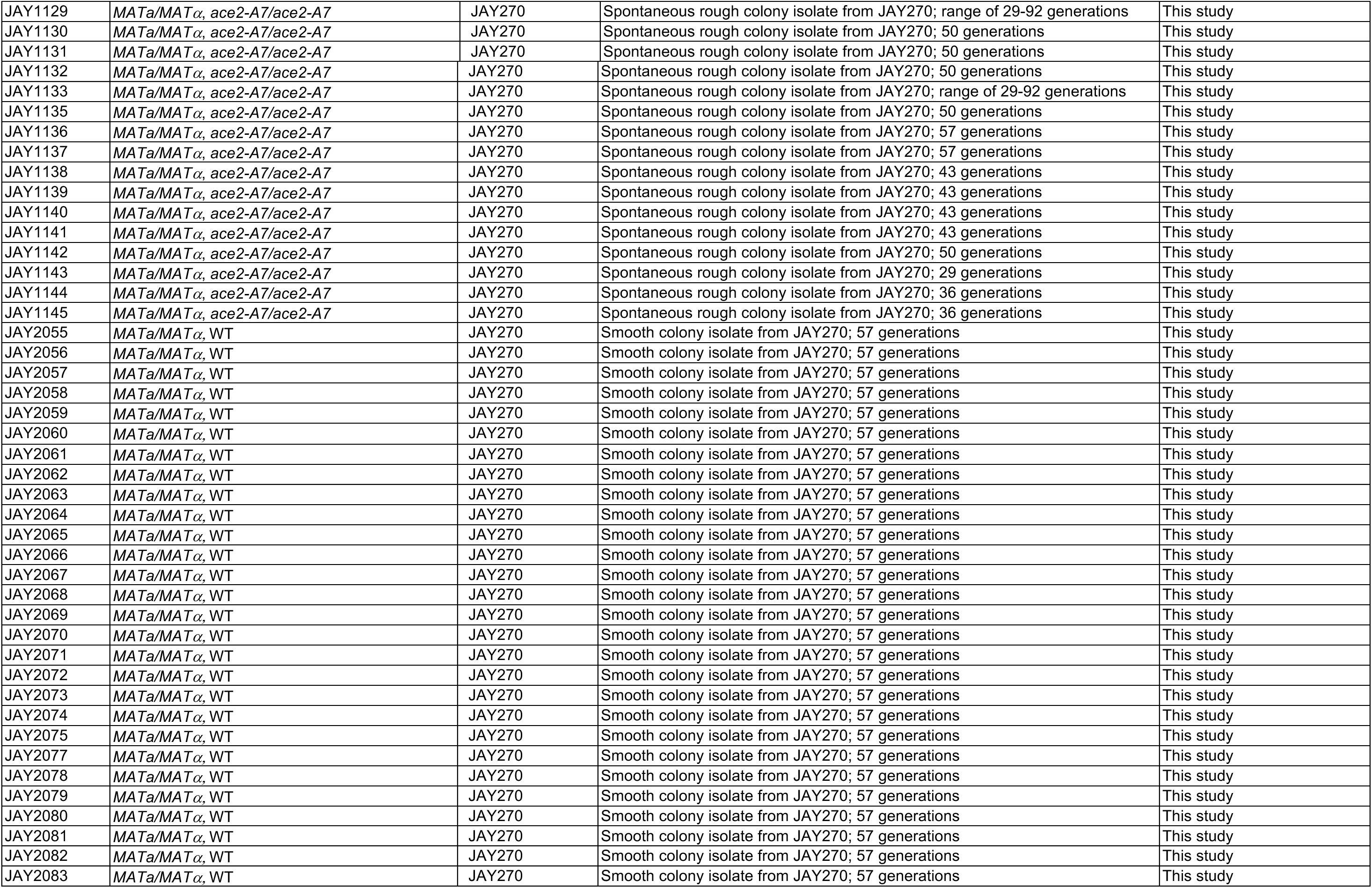
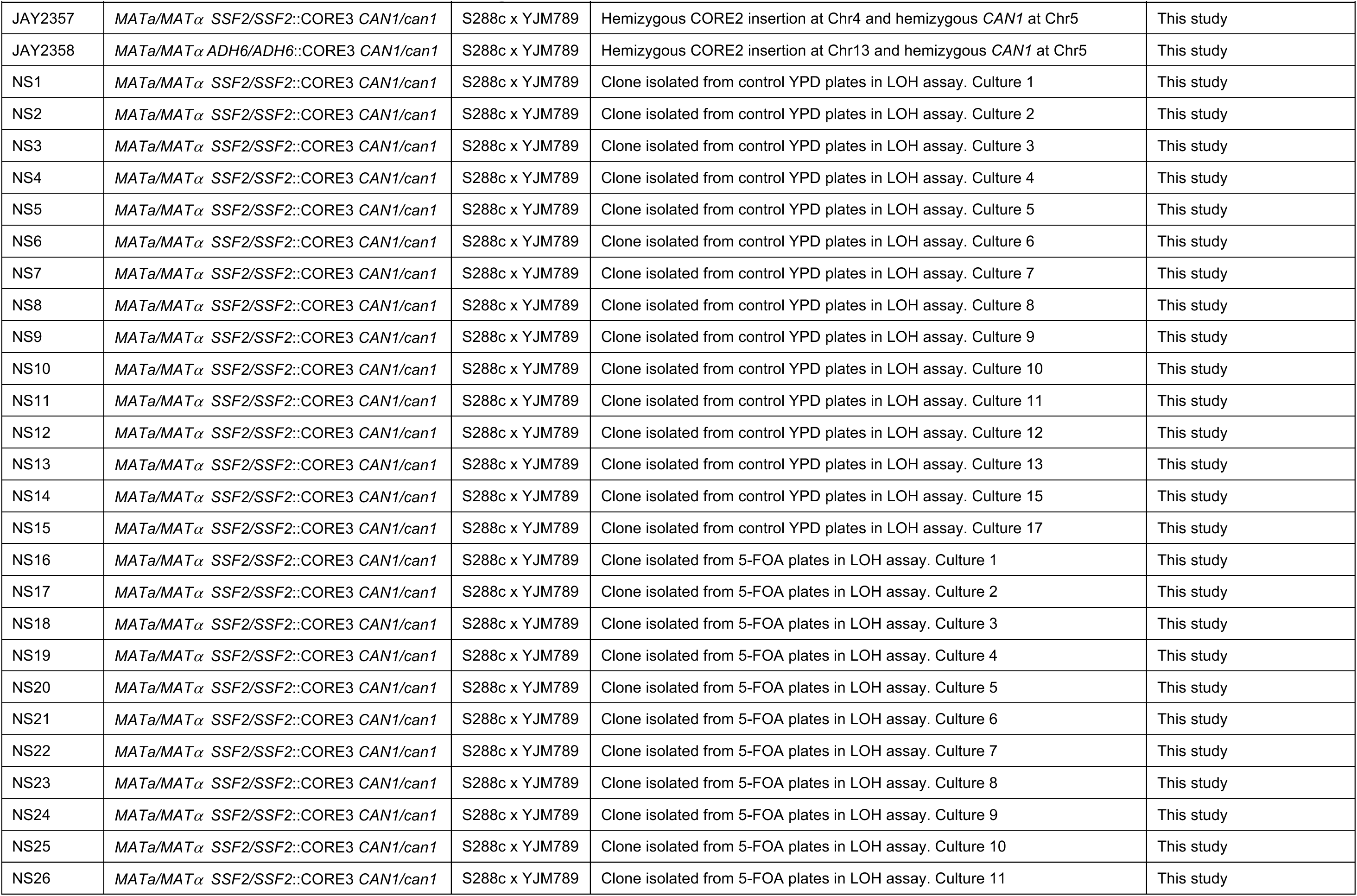
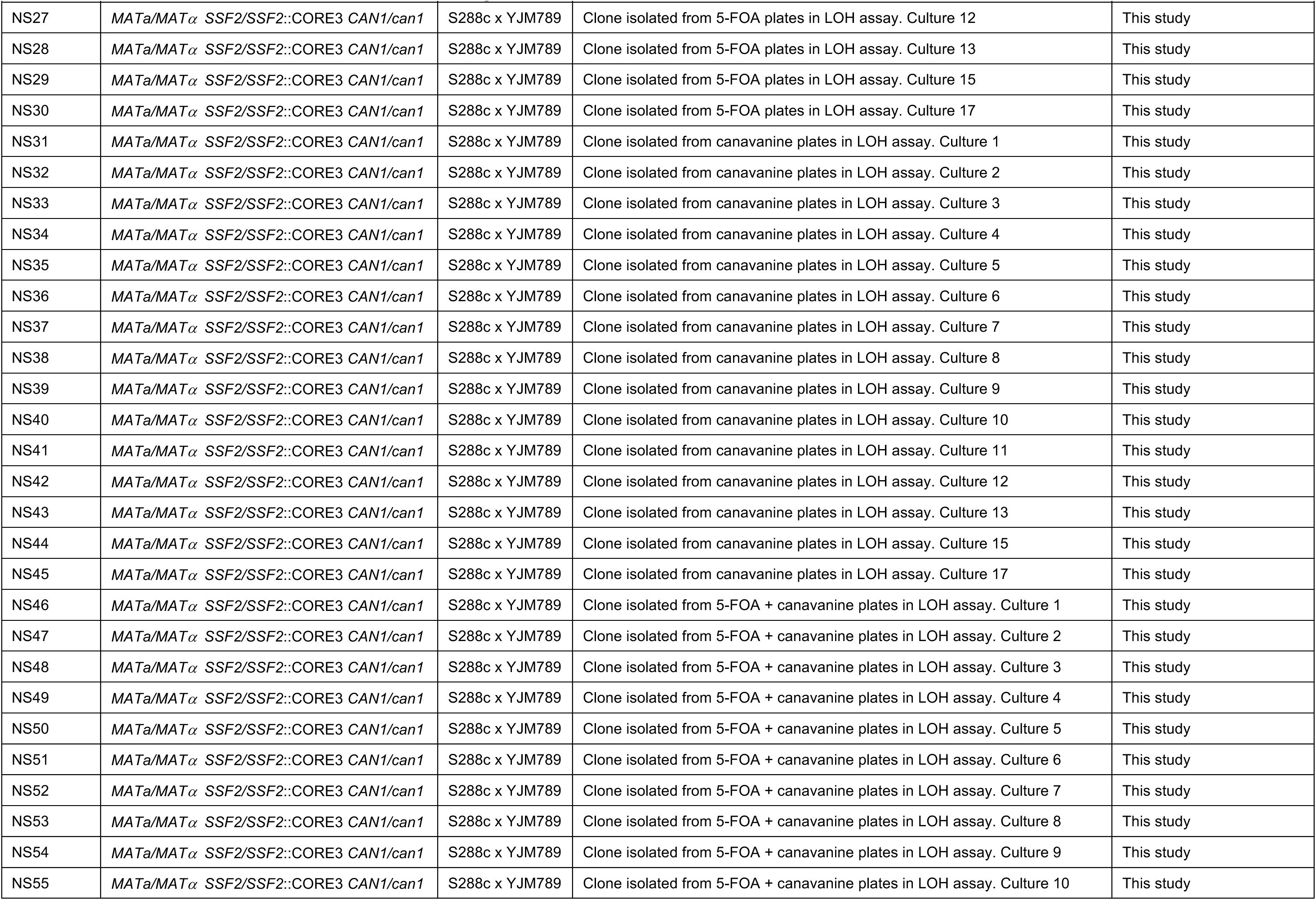
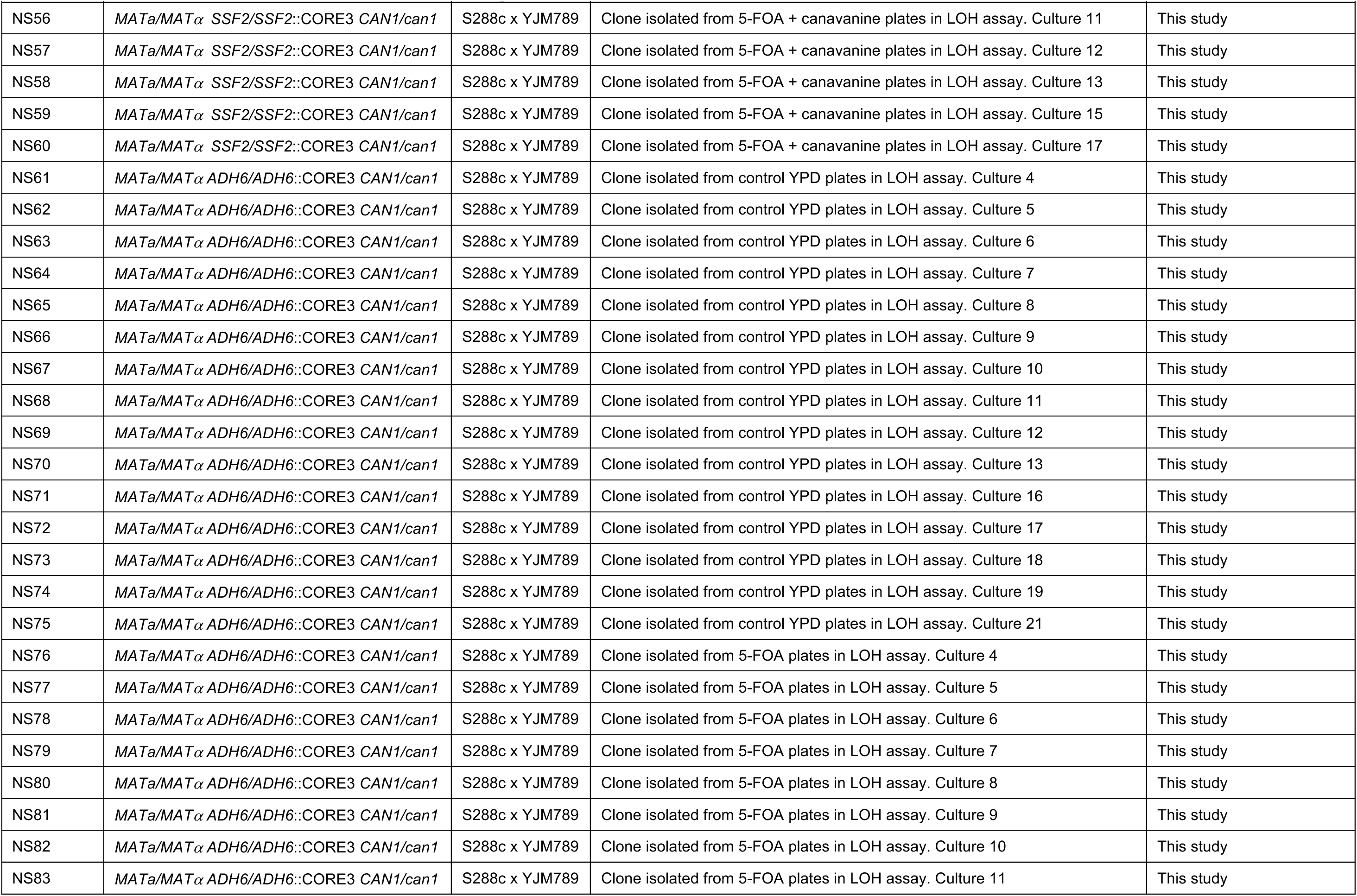
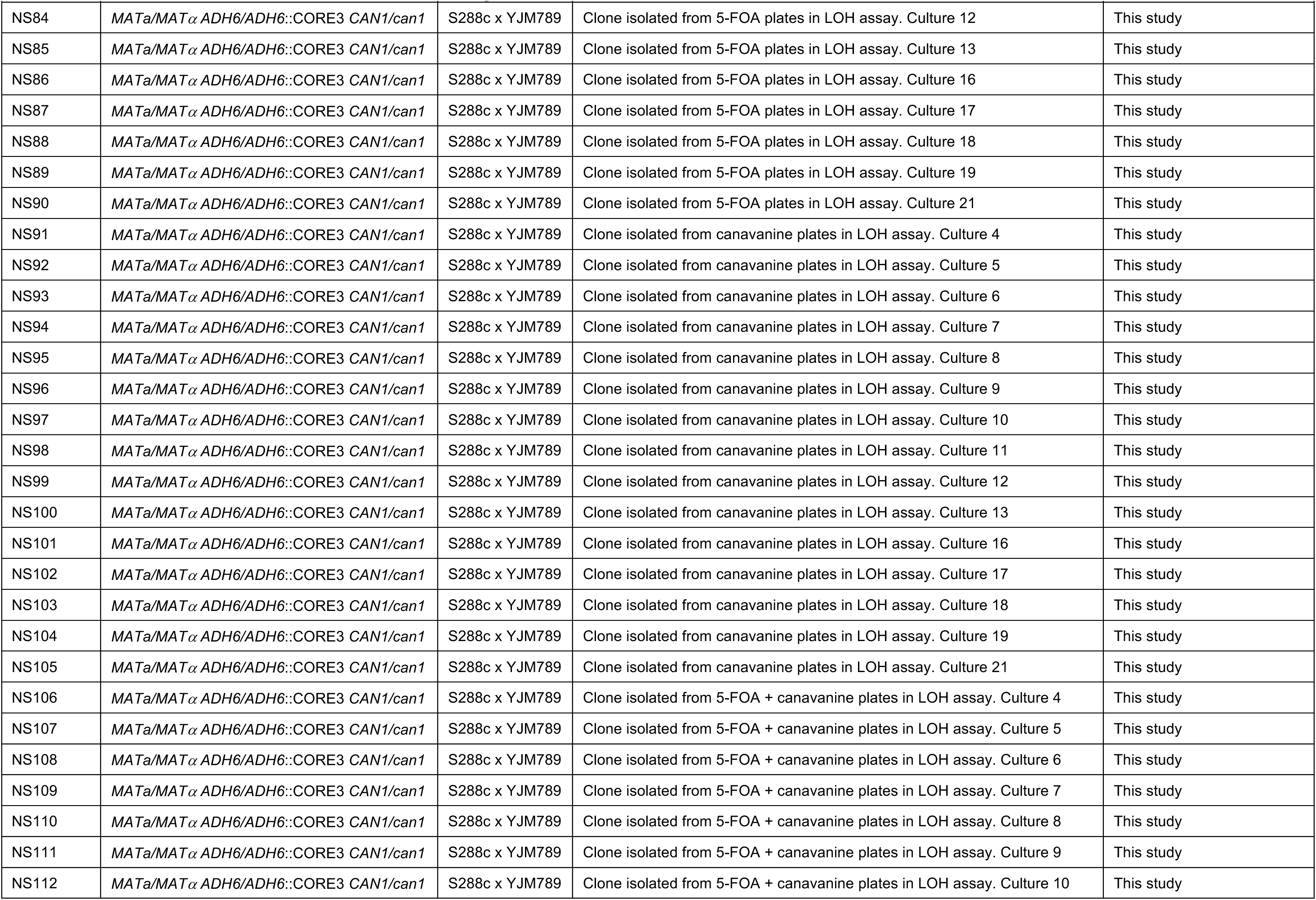
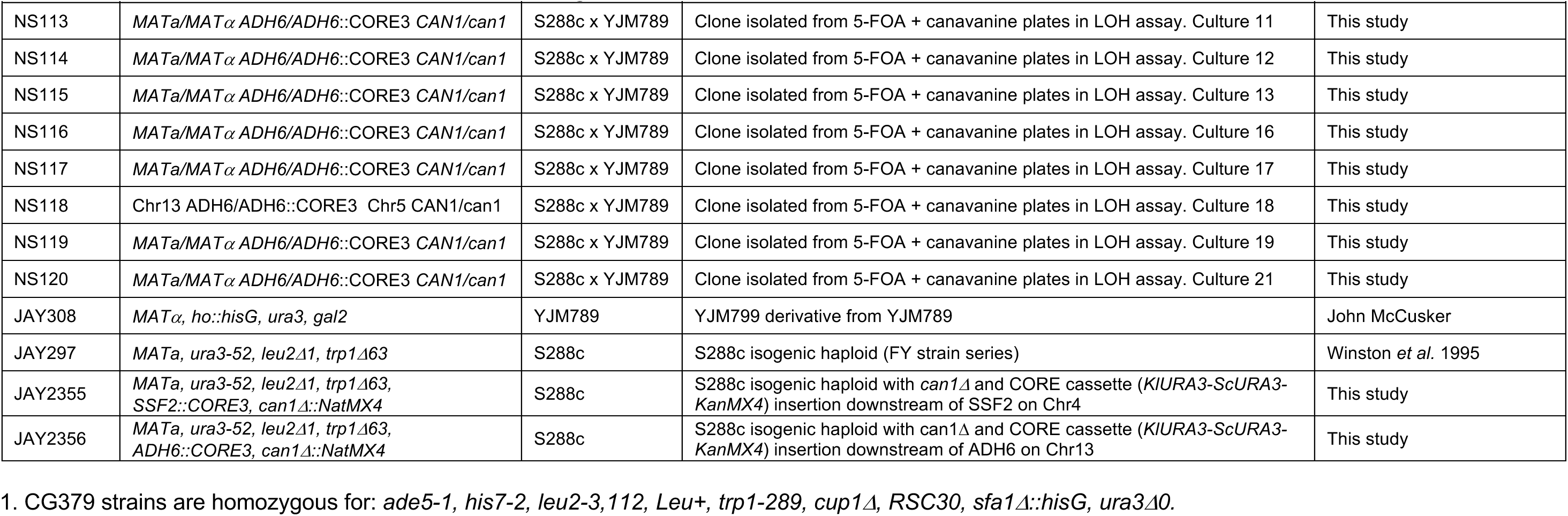
Yeast strains used in this study.

**Table S2.**
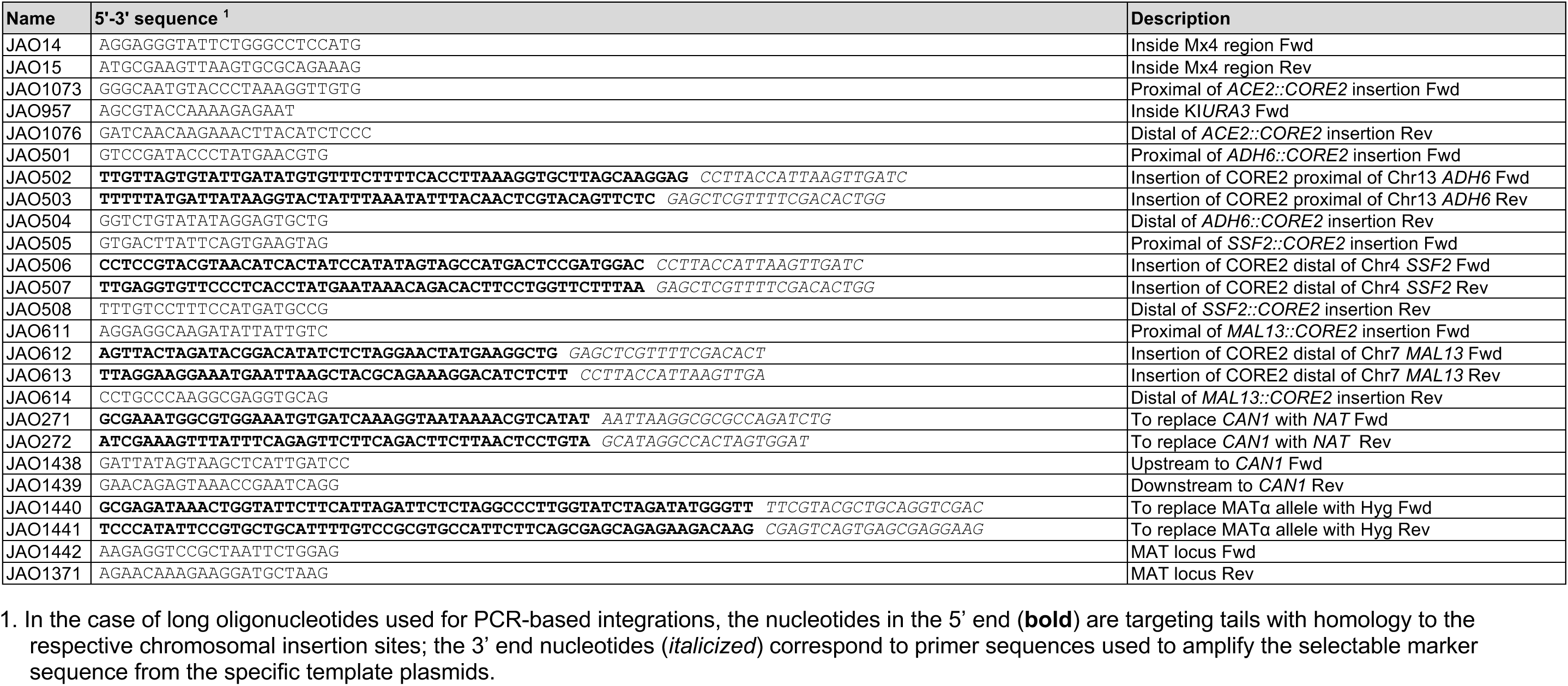
Oligonucleotides used in this study.

**Table S3.**
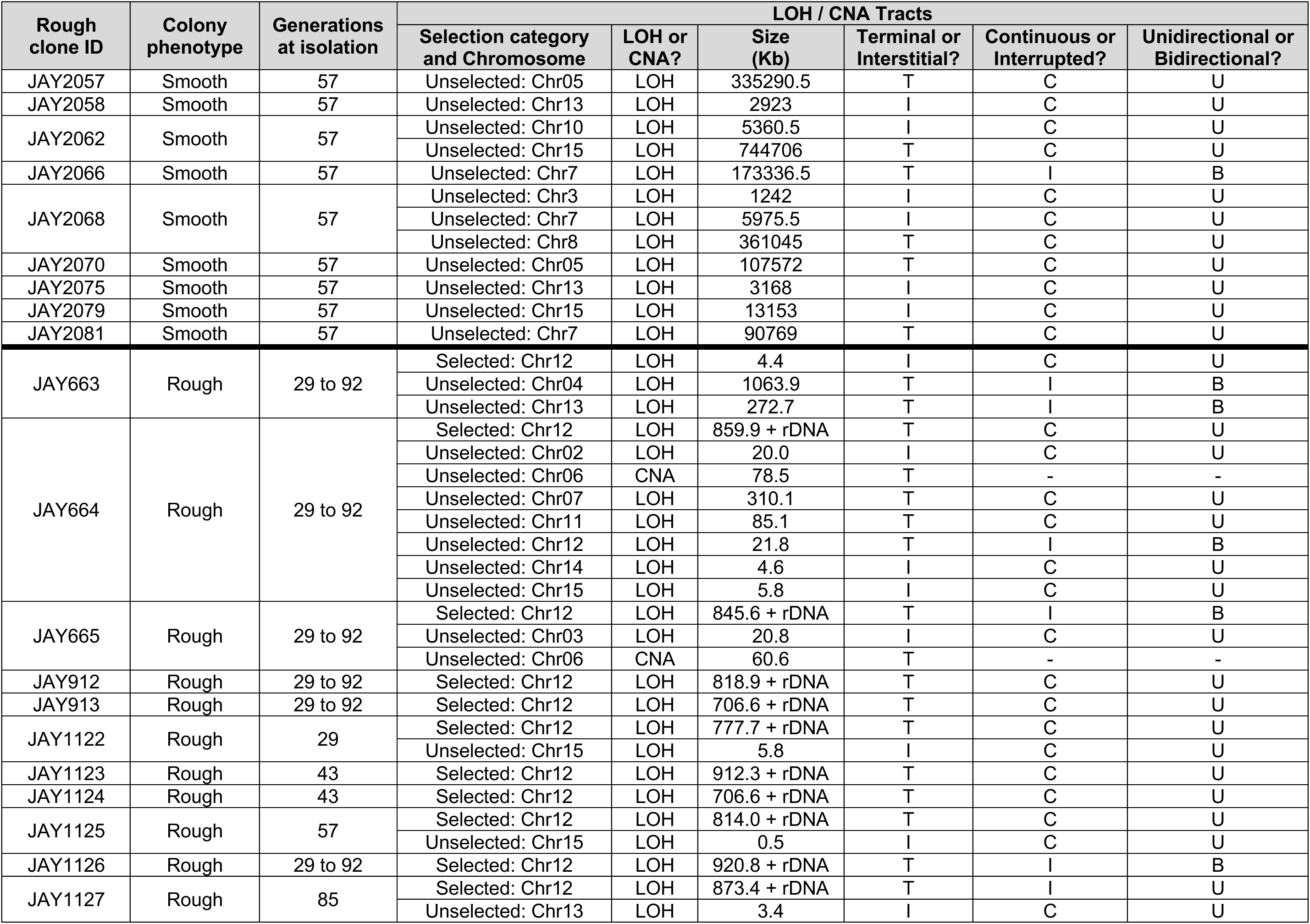
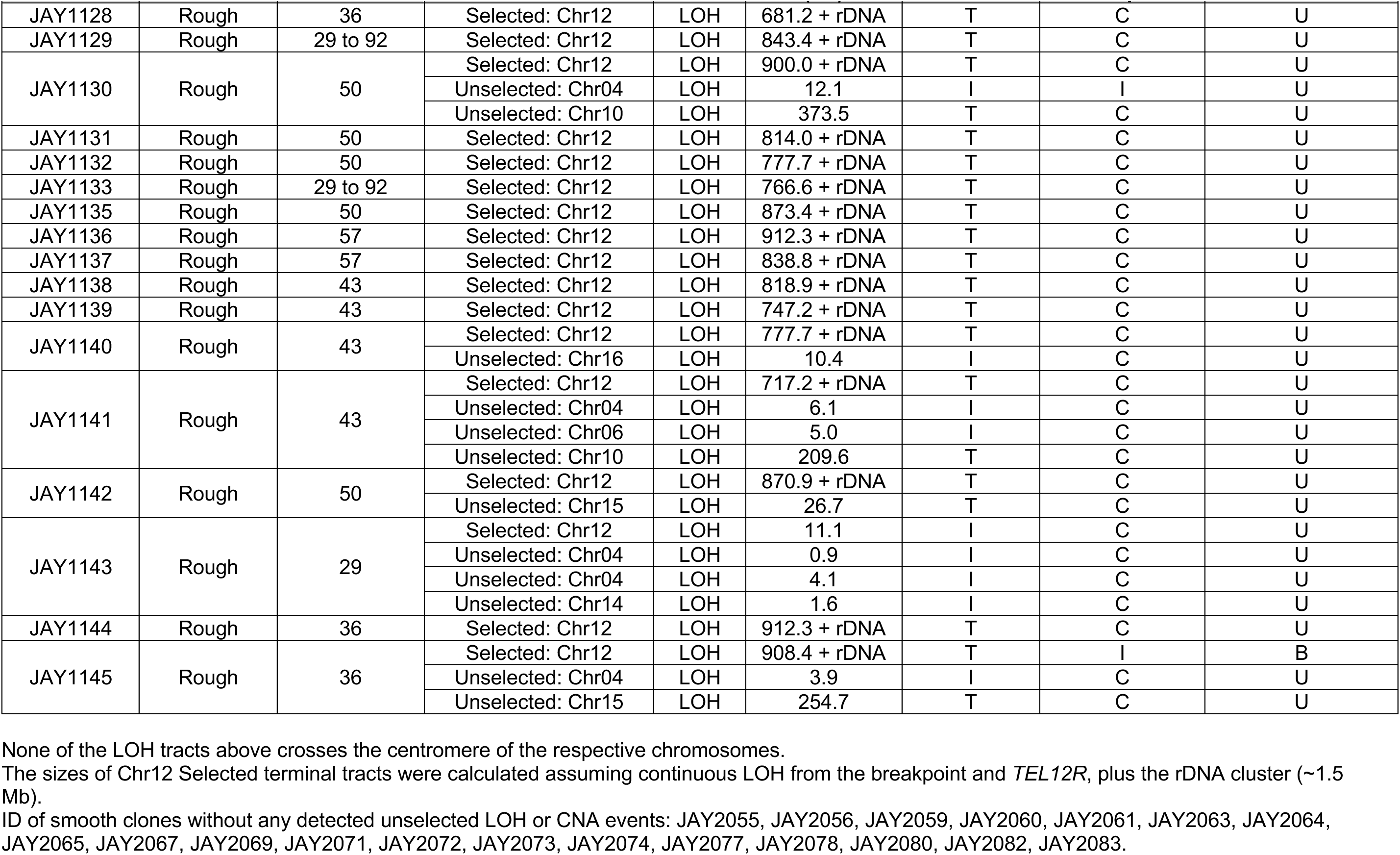
Summary of WGS analysis of smooth and rough colony isolates.

| Rough clone ID | Colony phenotype | Generations at isolation | LOH / CNA Tracts |  |  |  |  |  |
| --- | --- | --- | --- | --- | --- | --- | --- | --- |
|  |  |  | Selection category and Chromosome | LOH or CNA? | Size (Kb) | Terminal or Interstitial? | Continuous or Interrupted? | Unidirectional or Bidirectional? |
| JAY2057 | Smooth | 57 | Unselected: Chr05 | LOH | 335290.5 | T | C | U |
| JAY2058 | Smooth | 57 | Unselected: Chr13 | LOH | 2923 | I | C | U |
| JAY2062 | Smooth | 57 | Unselected: Chr10 | LOH | 5360.5 | I | C | U |
|  |  |  | Unselected: Chr15 | LOH | 744706 | T | C | U |
| JAY2066 | Smooth | 57 | Unselected: Chr7 | LOH | 173336.5 | T | I | B |
| JAY2068 | Smooth | 57 | Unselected: Chr3 | LOH | 1242 | I | C | U |
|  |  |  | Unselected: Chr7 | LOH | 5975.5 | I | C | U |
|  |  |  | Unselected: Chr8 | LOH | 361045 | T | C | U |
| JAY2070 | Smooth | 57 | Unselected: Chr05 | LOH | 107572 | T | C | U |
| JAY2075 | Smooth | 57 | Unselected: Chr13 | LOH | 3168 | I | C | U |
| JAY2079 | Smooth | 57 | Unselected: Chr15 | LOH | 13153 | I | C | U |
| JAY2081 | Smooth | 57 | Unselected: Chr7 | LOH | 90769 | T | C | U |
| JAY663 | Rough | 29 to 92 | Selected: Chr12 | LOH | 4.4 | I | C | U |
|  |  |  | Unselected: Chr04 | LOH | 1063.9 | T | I | B |
|  |  |  | Unselected: Chr13 | LOH | 272.7 | T | I | B |
| JAY664 | Rough | 29 to 92 | Selected: Chr12 | LOH | 859.9 + rDNA | T | C | U |
|  |  |  | Unselected: Chr02 | LOH | 20.0 | I | C | U |
|  |  |  | Unselected: Chr06 | CNA | 78.5 | T | - | - |
|  |  |  | Unselected: Chr07 | LOH | 310.1 | T | C | U |
|  |  |  | Unselected: Chr11 | LOH | 85.1 | T | C | U |
|  |  |  | Unselected: Chr12 | LOH | 21.8 | T | I | B |
|  |  |  | Unselected: Chr14 | LOH | 4.6 | I | C | U |
|  |  |  | Unselected: Chr15 | LOH | 5.8 | I | C | U |
| JAY665 | Rough | 29 to 92 | Selected: Chr12 | LOH | 845.6 + rDNA | T | I | B |
|  |  |  | Unselected: Chr03 | LOH | 20.8 | I | C | U |
|  |  |  | Unselected: Chr06 | CNA | 60.6 | T | - | - |
| JAY912 | Rough | 29 to 92 | Selected: Chr12 | LOH | 818.9 + rDNA | T | C | U |
| JAY913 | Rough | 29 to 92 | Selected: Chr12 | LOH | 706.6 + rDNA | T | C | U |
| JAY1122 | Rough | 29 | Selected: Chr12 | LOH | 777.7 + rDNA | T | C | U |
|  |  |  | Unselected: Chr15 | LOH | 5.8 | I | C | U |
| JAY1123 | Rough | 43 | Selected: Chr12 | LOH | 912.3 + rDNA | T | C | U |
| JAY1124 | Rough | 43 | Selected: Chr12 | LOH | 706.6 + rDNA | T | C | U |
| JAY1125 | Rough | 57 | Selected: Chr12 | LOH | 814.0 + rDNA | T | C | U |
|  |  |  | Unselected: Chr15 | LOH | 0.5 | I | C | U |
| JAY1126 | Rough | 29 to 92 | Selected: Chr12 | LOH | 920.8 + rDNA | T | I | B |
| JAY1127 | Rough | 85 | Selected: Chr12 | LOH | 873.4 + rDNA | T | I | U |
|  |  |  | Unselected: Chr13 | LOH | 3.4 | I | C | U |

**Table S3 (continued).** Summary of WGS analysis of smooth and rough colony isolates.
| Rough clone ID | Colony phenotype | Generations at isolation | LOH / CNA Tracts |  |  |  |  |  |
| --- | --- | --- | --- | --- | --- | --- | --- | --- |
|  |  |  | Selection category and Chromosome | LOH or CNA? | Size (Kb) | Terminal or Interstitial? | Continuous or Interrupted? | Unidirectional or Bidirectional? |
| JAY1128 | Rough | 36 | Selected: Chr12 | LOH | 681.2 + rDNA | T | C | U |
| JAY1129 | Rough | 29 to 92 | Selected: Chr12 | LOH | 843.4 + rDNA | T | C | U |
| JAY1130 | Rough | 50 | Selected: Chr12 | LOH | 900.0 + rDNA | T | C | U |
|  |  |  | Unselected: Chr04 | LOH | 12.1 | I | I | U |
|  |  |  | Unselected: Chr10 | LOH | 373.5 | T | C | U |
| JAY1131 | Rough | 50 | Selected: Chr12 | LOH | 814.0 + rDNA | T | C | U |
| JAY1132 | Rough | 50 | Selected: Chr12 | LOH | 777.7 + rDNA | T | C | U |
| JAY1133 | Rough | 29 to 92 | Selected: Chr12 | LOH | 766.6 + rDNA | T | C | U |
| JAY1135 | Rough | 50 | Selected: Chr12 | LOH | 873.4 + rDNA | T | C | U |
| JAY1136 | Rough | 57 | Selected: Chr12 | LOH | 912.3 + rDNA | T | C | U |
| JAY1137 | Rough | 57 | Selected: Chr12 | LOH | 838.8 + rDNA | T | C | U |
| JAY1138 | Rough | 43 | Selected: Chr12 | LOH | 818.9 + rDNA | T | C | U |
| JAY1139 | Rough | 43 | Selected: Chr12 | LOH | 747.2 + rDNA | T | C | U |
| JAY1140 | Rough | 43 | Selected: Chr12 | LOH | 777.7 + rDNA | T | C | U |
|  |  |  | Unselected: Chr16 | LOH | 10.4 | I | C | U |
| JAY1141 | Rough | 43 | Selected: Chr12 | LOH | 717.2 + rDNA | T | C | U |
|  |  |  | Unselected: Chr04 | LOH | 6.1 | I | C | U |
|  |  |  | Unselected: Chr06 | LOH | 5.0 | I | C | U |
|  |  |  | Unselected: Chr10 | LOH | 209.6 | T | C | U |
| JAY1142 | Rough | 50 | Selected: Chr12 | LOH | 870.9 + rDNA | T | C | U |
|  |  |  | Unselected: Chr15 | LOH | 26.7 | T | C | U |
| JAY1143 | Rough | 29 | Selected: Chr12 | LOH | 11.1 | I | C | U |
|  |  |  | Unselected: Chr04 | LOH | 0.9 | I | C | U |
|  |  |  | Unselected: Chr04 | LOH | 4.1 | I | C | U |
|  |  |  | Unselected: Chr14 | LOH | 1.6 | I | C | U |
| JAY1144 | Rough | 36 | Selected: Chr12 | LOH | 912.3 + rDNA | T | C | U |
| JAY1145 | Rough | 36 | Selected: Chr12 | LOH | 908.4 + rDNA | T | I | B |
|  |  |  | Unselected: Chr04 | LOH | 3.9 | I | C | U |
|  |  |  | Unselected: Chr15 | LOH | 254.7 | T | C | U |

None of the LOH tracts above crosses the centromere of the respective chromosomes.
The sizes of Chr12 Selected terminal tracts were calculated assuming continuous LOH from the breakpoint and *TEL12R*, plus the rDNA cluster (~1.5 Mb).
ID of smooth clones without any detected unselected LOH or CNA events: JAY2055, JAY2056, JAY2059, JAY2060, JAY2061, JAY2063, JAY2064, JAY2065, JAY2067, JAY2069, JAY2071, JAY2072, JAY2073, JAY2074, JAY2077, JAY2078, JAY2080, JAY2082, JAY2083.

**Table S4.** Summary of LOH rate analyses.

| Chromosome<sup>a</sup> | Observed or Expected<sup>b</sup> | Cultures<sup>c</sup> (Volume) | Mutation Rate | Lower 95% C.I. | Upper 95% C.I. | Obs/Exp ratio |
| --- | --- | --- | --- | --- | --- | --- |
| Figure 3A: JAY270 strain background, *MATa/MATα* ; L&C method of the median |  |  |  |  |  |  |
| Chr4 | Obs | 20 (5 ml) | 3.47E-05 | 1.73E-05 | 5.65E-05 | - |
| Chr13 | Obs | 20 (5 ml) | 4.02E-05 | 2.61E-05 | 5.84E-05 | - |
| Chr5 | Obs | 63 (5 ml) | 1.06E-04 | 9.33E-05 | 1.15E-04 | - |
| Chr4 + Chr5 | Exp | - | 3.67E-09 | 1.62E-09 | 6.48E-09 | 27 |
| Chr4 + Chr5 | Obs | 21 (5 ml) | 9.76E-08 | 3.52E-08 | 1.32E-07 |  |
| Chr13 + Chr5 | Exp | - | 4.24E-09 | 2.44E-09 | 6.70E-09 | 14 |
| Chr13 + Chr5 | Obs | 21 (5 ml) | 6.14E-08 | - | 1.23E-07 |  |
| Figure 3B: CG379 strain background, *MATa/MATα* ; L&C method of the median |  |  |  |  |  |  |
| Chr4 | Obs | 23 (5 ml) | 1.26E-04 | 8.56E-05 | 1.75E-04 | - |
| Chr13 | Obs | 25 (5 ml) | 7.30E-05 | 4.71E-05 | 1.16E-04 | - |
| Chr5 | Obs | 41 (5 ml) | 7.10E-05 | 6.62E-05 | 8.02E-05 | - |
| Chr4 + Chr5 | Exp | - | 8.96E-09 | 5.67E-09 | 1.40E-08 | 30 |
| Chr4 + Chr5 | Obs | 22 (5 ml) | 2.65E-07 | 2.29E-07 | 5.18E-07 |  |
| Chr13 + Chr5 | Exp | - | 5.19E-09 | 3.12E-09 | 9.31E-09 | 55 |
| Chr13 + Chr5 | Obs | 21 (5 ml) | 2.86E-07 | 2.00E-07 | 5.19E-07 |  |
| Figure 3B: CG379 strain background, *MATa/matΔ* ; L&C method of the median |  |  |  |  |  |  |
| Chr4 | Obs | 20 (5 ml) | 4.90E-05 | 4.50E-05 | 6.90E-05 | - |
| Chr13 | Obs | 21 (5 ml) | 3.55E-05 | 2.99E-05 | 4.30E-05 | - |
| Chr5 | Obs | 21 (5 ml) | 5.65E-05 | 4.21E-05 | 7.32E-05 | - |
| Chr4 + Chr5 | Exp | - | 2.77E-09 | 1.89E-09 | 5.05E-09 | 50 |
| Chr4 + Chr5 | Obs | 21 (5 ml) | 1.39E-07 | 7.45E-08 | 2.28E-07 |  |
| Chr13 + Chr5 | Exp | - | 2.00E-09 | 1.26E-09 | 3.15E-09 | 51 |
| Chr13 + Chr5 | Obs | 20 (5 ml) | 1.03E-07 | 7.26E-08 | 1.55E-07 |  |
| Figure 3C: S288c x YJM789 strain background, *MATa/MATα* ; L&C method of the median |  |  |  |  |  |  |
| Chr4 | Obs | 21 (5 ml) | 1.65E-05 | 1.38E-05 | 2.09E-05 | - |
| Chr13 | Obs | 21 (5 ml) | 1.86E-05 | 1.33E-05 | 2.12E-05 | - |
| Chr5 | Obs | 41 (5 ml) | 4.00E-05 | 3.47E-05 | 4.59E-05 | - |
| Chr4-Chr5 | Exp | - | 6.58E-10 | 4.77E-10 | 9.61E-10 | 66 and 70 |
| Chr4 + Chr5 | Exp matched | 20 x 20 (5 ml) | 6.21E-10 | 5.03E-10 | 7.56E-10 |  |
| Chr4 + Chr5 | Obs | 21 (5 ml) | 4.32E-08 | 2.40E-08 | 5.53E-08 |  |
| Chr13 + Chr5 | Exp | - | 7.45E-10 | 4.62E-10 | 9.73E-10 | 28 and 27 |
| Chr13 + Chr5 | Exp matched | 21 x 21 (5 ml) | 7.58E-10 | 5.33E-10 | 1.05E-09 |  |
| Chr13 + Chr5 | Obs | 21 (5 ml) | 2.06E-08 | 1.43E-08 | 3.15E-08 |  |

**Table S4 (continued).** Summary of LOH rate analyses.
| Chromosome<sup>a</sup> | Observed or Expected<sup>b</sup> | Cultures<sup>c</sup> (Volume) | Mutation Rate | Lower 95% C.I. | Upper 95% C.I. | Obs/Exp ratio |
| --- | --- | --- | --- | --- | --- | --- |
| Figure S7A: S288c x YJM789 strain background, *MATa/MATα* ; L&C method of the median |  |  |  |  |  |  |
| Chr4 | Obs | 15 (20 μl) | 2.01E-05 | 1.27E-05 | 2.99E-05 | - |
| Chr13 | Obs | 15 (20 μl) | 3.64E-05 | 9.00E-06 | 5.04E-05 | - |
| Chr5 | Obs | 30 (20 μl) | 3.57E-05 | 2.71E-05 | 4.50E-05 | - |
| Chr4 + Chr5 | Exp | - | 7.18E-10 | 3.43E-10 | 1.35E-09 | 60 |
| Chr4 + Chr5 | Obs | 21 (5 ml) | 4.32E-08 | 2.40E-08 | 5.53E-08 |  |
| Chr13 + Chr5 | Exp | - | 1.30E-09 | 2.44E-10 | 2.27E-09 | 16 |
| Chr13 + Chr5 | Obs | 21 (5 ml) | 2.06E-08 | 1.43E-08 | 3.15E-08 |  |
| Figure S7B: S288c x YJM789 strain background, *MATa/MATα* ; MSS – Maximum Likelihood Estimate |  |  |  |  |  |  |
| Chr4 | Obs | 21 (5 ml) | 5.93E-06 | 4.63E-06 | 7.36E-06 | - |
| Chr13 | Obs | 21 (5 ml) | 6.24E-06 | 4.86E-06 | 7.74E-06 | - |
| Chr5 | Obs | 41 (5 ml) | 1.32E-05 | 1.15E-05 | 1.49E-05 | - |
| Chr4 + Chr5 | Exp | - | 7.82E-11 | 5.33E-11 | 1.10E-10 | 150 |
| Chr4 + Chr5 | Obs | 21 (5 ml) | 1.17E-08 | 6.90E-09 | 1.74E-08 |  |
| Chr13 + Chr5 | Exp | - | 8.22E-11 | 5.59E-11 | 1.15E-10 | 129 |
| Chr13 + Chr5 | Obs | 21 (5 ml) | 1.06E-08 | 6.08E-09 | 1.61E-08 |  |
| Figure S7C: S288c x YJM789 strain background, *MATa/MATα* ; MSS – Maximum Likelihood Estimate |  |  |  |  |  |  |
| Chr4 | Obs | 15 (20 μl) | 1.51E-05 | 1.13E-05 | 1.92E-05 | - |
| Chr13 | Obs | 15 (20 μl) | 1.07E-05 | 7.66E-06 | 1.41E-05 | - |
| Chr5 | Obs | 30 (20 μl) | 3.26E-05 | 2.75E-05 | 3.81E-05 | - |
| Chr4 + Chr5 | Exp | - | 4.92E-10 | 3.11E-10 | 7.33E-10 | 24 |
| Chr4 + Chr5 | Obs | 21 (5 ml) | 1.17E-08 | 6.90E-09 | 1.74E-08 |  |
| Chr13 + Chr5 | Exp | - | 3.50E-10 | 2.11E-10 | 5.38E-10 | 30 |
| Chr13 + Chr5 | Obs | 21 (5 ml) | 1.06E-08 | 6.08E-09 | 1.61E-08 |  |

a. Indicates the genomic region of selection of either single or double LOH events.
b. Indicates whether the LOH rate displayed in each row is derived either from observed experimental data (Obs), or expected double LOH (Exp) based on independence calculated by multiplication of the two respective observed single LOH rates. Exp matched indicates expected double LOH rates calculated by multiplying the two individual observed single LOH rates within each specific matched culture.
c. Indicates the number of replicate cell cultures used in each rate measurement and the volume of liquid YPD they were grown in.
d. Mutation rates and 95% confidence intervals (C.I.) are given in LOH events per cell per division.

**Table S5.**
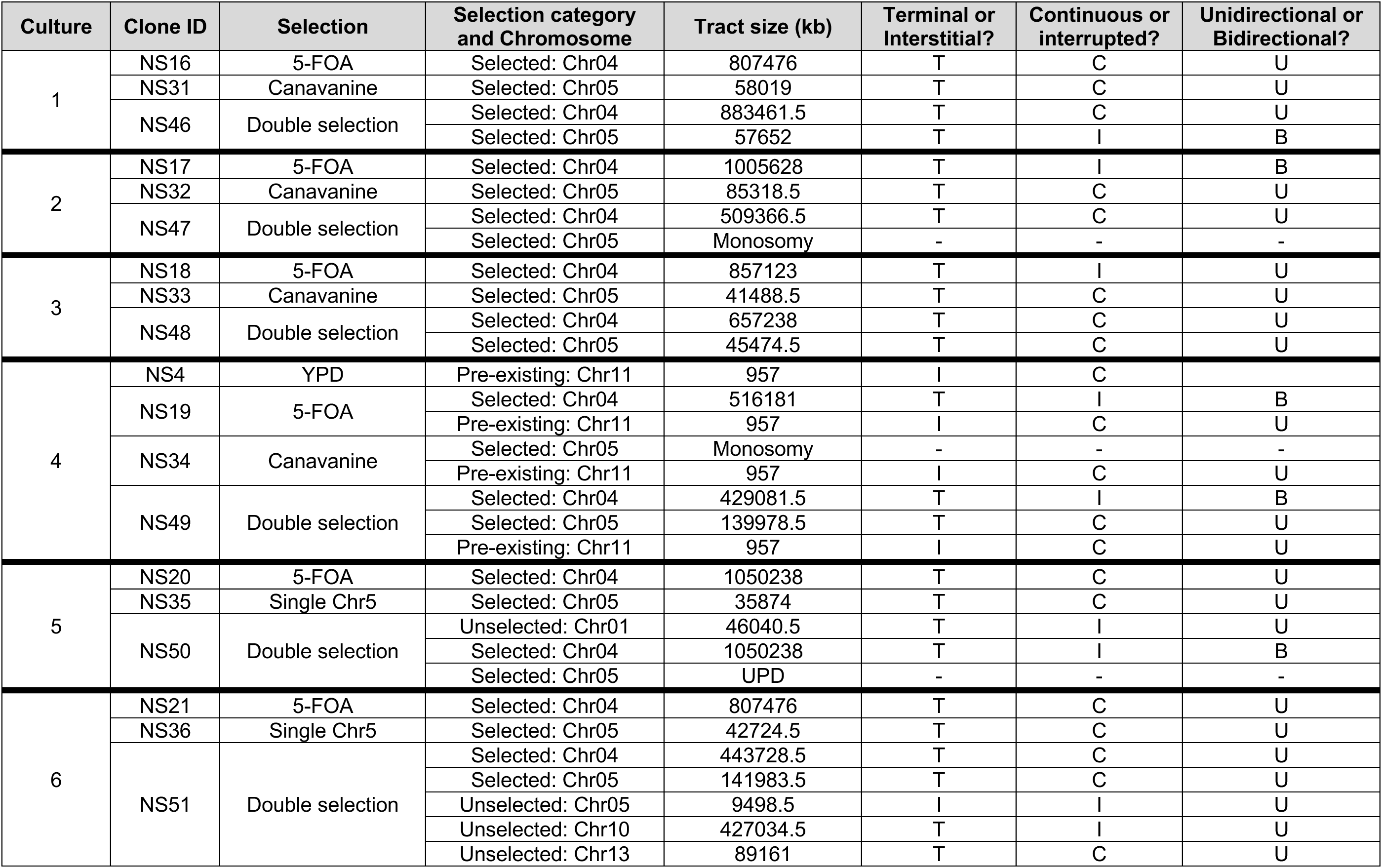
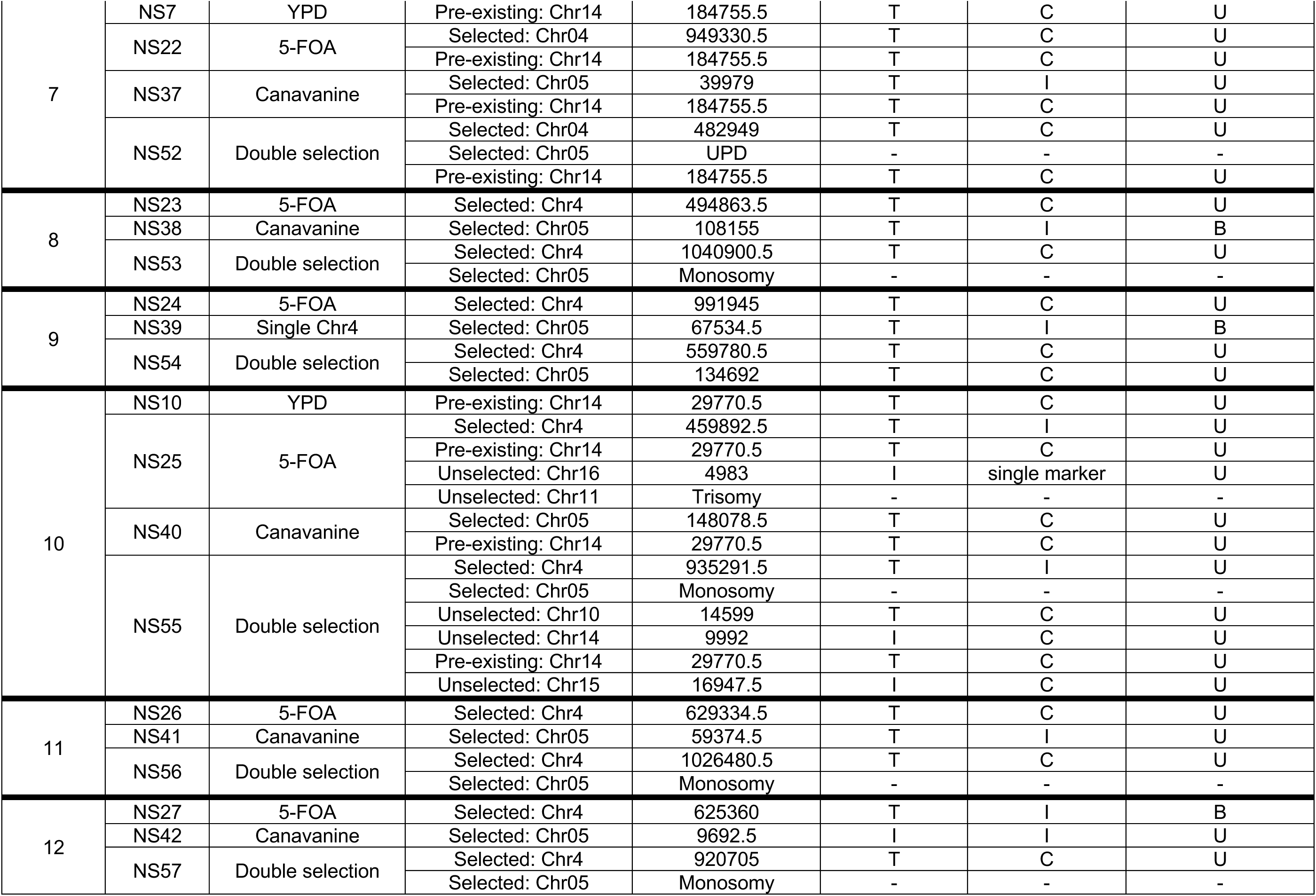
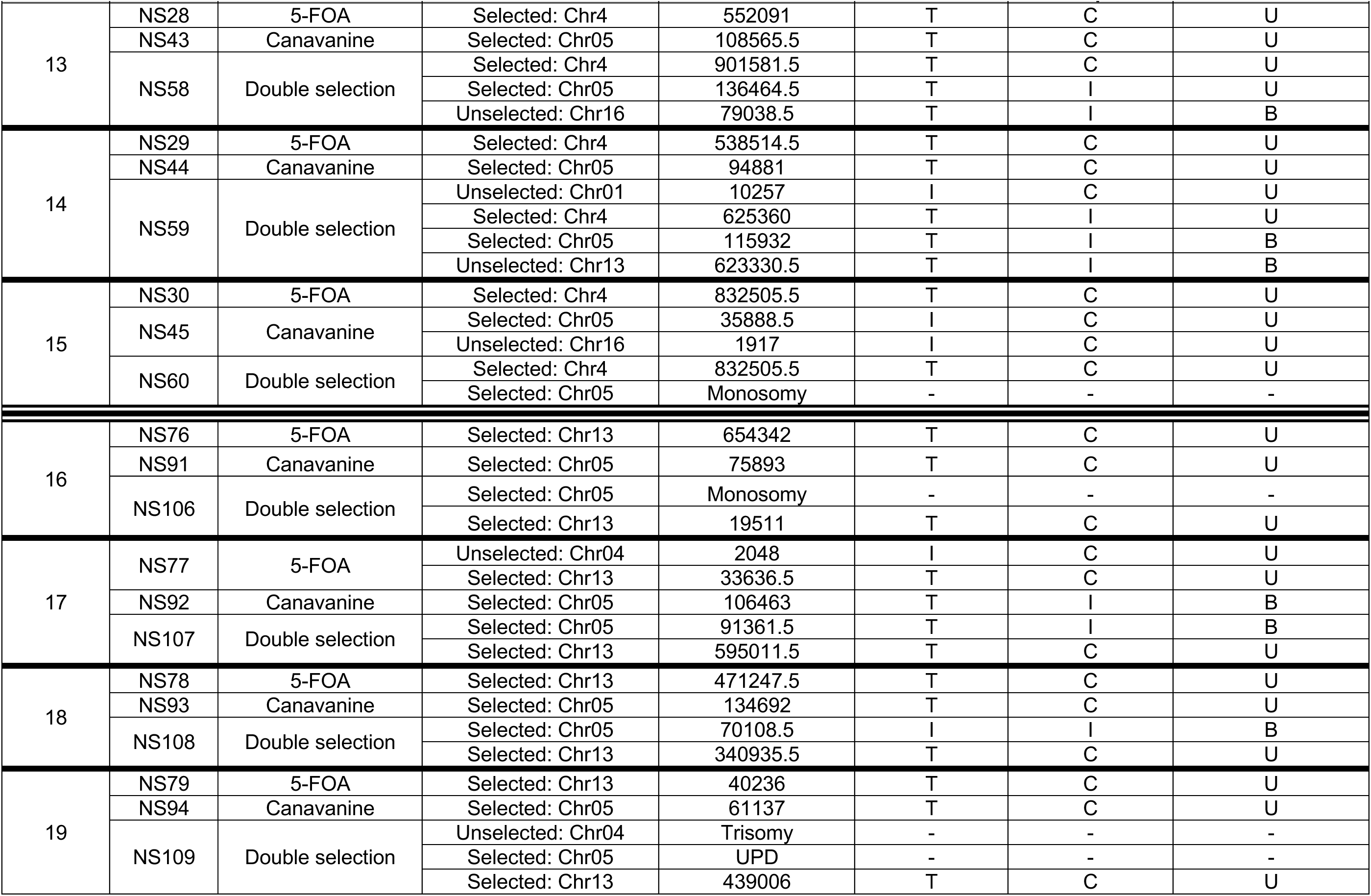
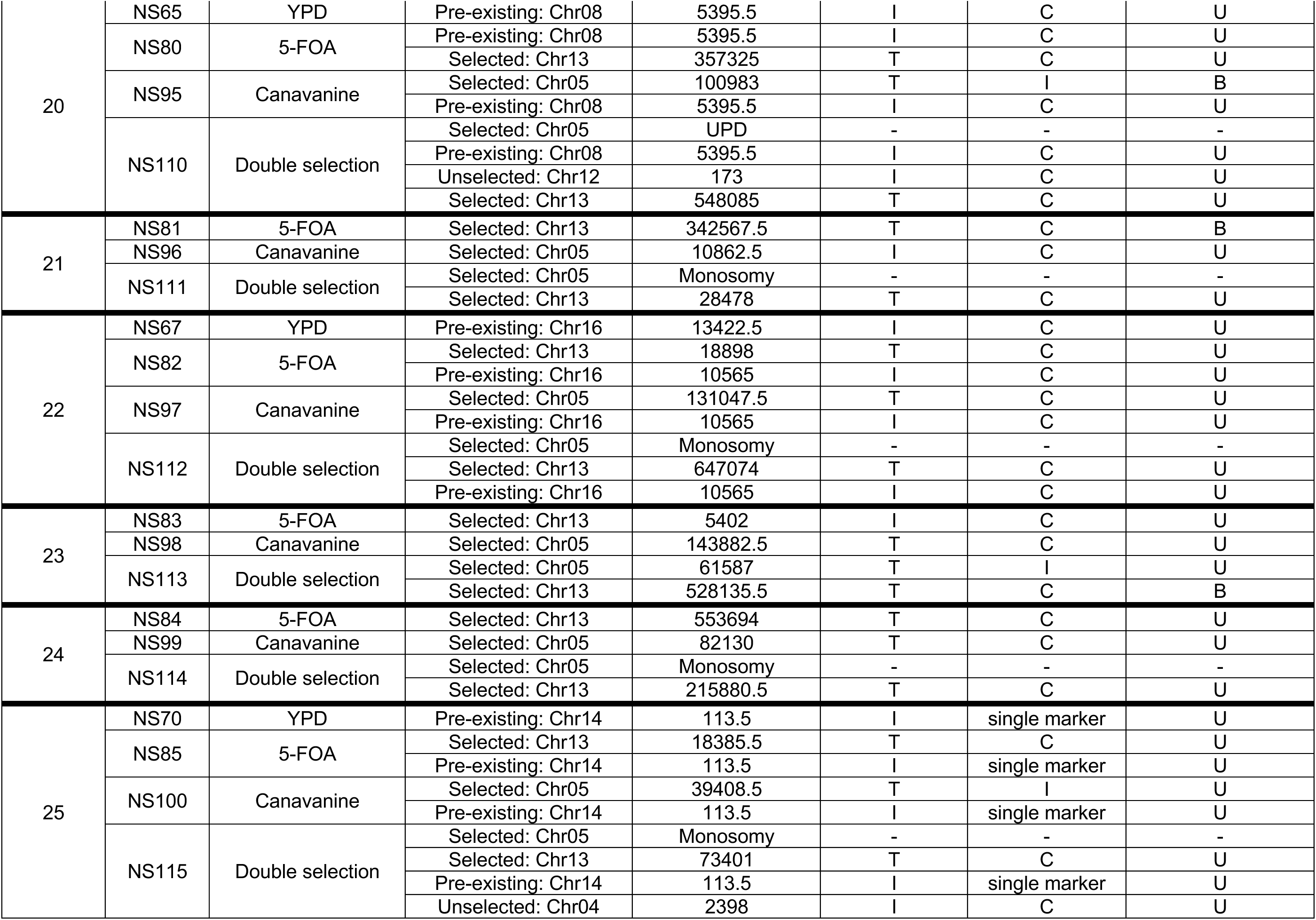
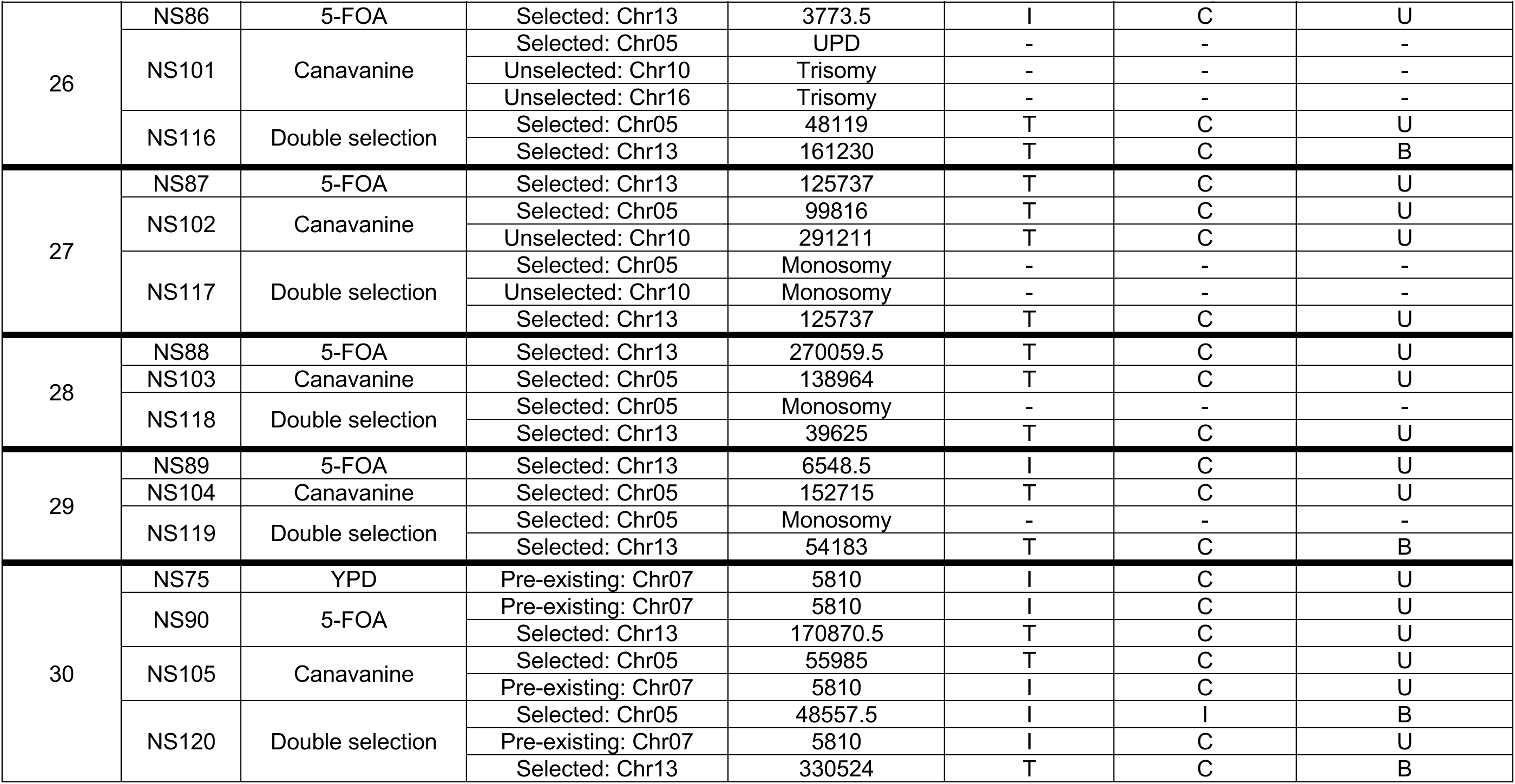
Summary of WGS analysis of S288c x YJM789 hybrid isolates.

